# APOE-NOTCH Axis Governs Elastogenesis During Human Cardiac Valve Remodeling

**DOI:** 10.1101/2023.04.26.538443

**Authors:** Ziyi Liu, Yu Liu, Zhiyun Yu, Nicole Pek, Anna O’Donnell, Ian Glass, David S. Winlaw, Minzhe Guo, Ya-Wen Chen, Joseph C. Wu, Katherine E. Yutzey, Yifei Miao, Mingxia Gu

## Abstract

**Background:** Valve remodeling is a complex process involving extracellular matrix organization, development of trilaminar structures, and physical elongation of valve leaflets. However, the cellular and molecular mechanisms regulating valve remodeling and their roles in congenital valve disorders remain poorly understood.

**Methods:** Semilunar valves and atrioventricular valves from healthy and age-matched human fetal hearts with pulmonary stenosis (PS) were collected. Single-Cell RNA-sequencing (scRNA-seq) was performed to determine the transcriptomic landscape of multiple valvular cell subtypes in valve remodeling and disease. Spatial localization of newly-identified cell subtypes was determined via immunofluorescence and RNA *in situ* hybridization. The molecular mechanisms mediating valve development was investigated utilizing primary human fetal heart valve interstitial cells (VICs) and endothelial cells (VECs).

**Results:** scRNA-seq analysis of healthy human fetal valves identified a novel APOE^+^ elastin-producing VIC subtype (Elastin-VICs) spatially located underneath VECs sensing the unidirectional flow. Knockdown of *APOE* in fetal VICs resulted in significant elastogenesis defects. In pulmonary valve with PS, we observed decreased expression of *APOE* and other genes regulating elastogenesis such as *EMILIN1* and *LOXL1*, as well as elastin fragmentation. These findings suggested the crucial role of APOE in regulating elastogenesis during valve remodeling. Furthermore, cell-cell interaction analysis revealed that JAG1 from unidirectional VECs activates NOTCH signaling in Elastin-VICs through NOTCH3. *In vitro* Jag1 treatment in VICs increased elastogenesis, while similar observations were found in VICs co-cultured with VECs in the presence of unidirectional flow. Notably, we found that the JAG1-NOTCH3 signaling pair was drastically reduced in the PS valves. Lastly, we demonstrated that APOE is indispensable for JAG1-induced NOTCH activation in VICs, reinforcing the presence of a synergistic intrinsic and external regulatory network involving APOE and NOTCH signaling that is responsible for regulating elastogenesis during human valve remodeling.

**Conclusion:** scRNA-seq analysis of human fetal valves identified a novel Elastin-VIC subpopulation, and revealed mechanism of intrinsic APOE and external NOTCH signaling in regulating elastogenesis during cardiac valve remodeling. These mechanisms may contribute to deciphering the pathogenesis of elastin malformation in congenital valve diseases.

**Clinical Perspective:** *What Is New?:*

- High-resolution single-cell transcriptome atlas generated from healthy human fetal heart valves and valves affected by pulmonary stenosis during the early phase of valve remodeling prior to birth.
- A unique subset of valve interstitial cells (VICs) that produce elastin (Elastin-VICs) was identified.
- Elastin-VICs specifically located underneath the valve endothelial cells (VECs) sensing unidirectional flow, and played a crucial role in elastin maturation via the expression of APOE.
- Elastin-VICs communicated with adjacent VECs via the JAG1-NOTCH signaling, facilitating elastin formation and valve remodeling.

*What Are the Clinical Implications?:*

- Elastin-VICs from patient valvular tissues with Pulmonary Stenosis exhibit decreased APOE-NOTCH signaling and elastin fragmentation.
- Direct targeting of APOE and NOTCH signaling could be a novel approach to promote elastin fiber formation and valve remodeling in patients with valvular defects.

## Introduction

During human embryonic and postnatal heart development, cardiac valve remodeling is vital for valvulogenesis. Valvular remodeling involves the physical elongation of valve leaflets, accompanied by an increase in extracellular matrix (ECM) production necessary for the development of the trilaminar valve structures – fibrosa, spongiosa, and ventricularis (semilunar valves) or atrialis (atrioventricular valves), which mainly comprise of collagen, glycosaminoglycan (GAG), and elastin respectively^1, 2^. At 14 weeks of gestation, valves are primarily composed of GAG after extensive Endocardial-Mesenchymal Transition (Endo-MT) in the atrioventricular canal and outflow tract^3^. By 36 weeks of gestation, the valves are stratified into three ECM layers which grow and mature with age, ultimately developing into distinct tri-laminar structures^1^. Proper valvular remodeling is critical for normal valve development; disruptions to the valvular remodeling process have been found to underlie valve diseases such as valve prolapse presented in Marfan syndrome, pulmonary valve stenosis in Tetralogy of Fallot (TOF), and aortic valve atresia in hypoplastic left heart syndrome (HLHS)^2, 4–6^.

To date, the cellular and molecular mechanisms driving valve remodeling during human heart development remain poorly understood, mainly due to the lack of equivalent prenatal valve remodeling period in rodents, and the limited accessibility and technical difficulties in getting human valve specimens during the later gestation and prenatal stages^7^. Key players in valve include valve interstitial cells (VICs), which are the predominant ECM-producing cells. VICs are responsible for the valve compartmentalization by adapting to dynamic mechanical forces and secreting different ECM components during early valve remodeling, although direct evidence is still limited^8^. In addition, early studies using animal models found that blood flow-induced shear stress regulates valve remodeling. Rodent studies have demonstrated the pivotal role of the endothelium/endocardium in sensing blood flow-induced shear stress and regulating WNT-dependent endocardial cushion remodeling^9^. Mechanical alteration to blood flow in chick and zebrafish resulted in valve malformation, leading to severe cardiac defects^10–13^. However, the exact mechanisms of unidirectional (laminar) flow and turbulent (oscillatory) flow in regulating VIC physiology, and therefore valve remodeling and ECM organization have not been well-established.

Recent studies have employed scRNA-seq and spatial transcriptomics to dissect the cellular landscape of valves in both humans and mice^14–16^. In humans, although multiple valvular cell types including VICs and valvular endothelial cells (VECs) have been revealed, VIC and VEC subtypes in human fetal valves subjected to early remodeling cues were not adequately characterized^14, 15^. In postnatal mouse valves, a subpopulation of collagen-producing VICs was found to reside adjacent to *Prox1*^+^ VECs that were in the direction of turbulent flow, while a subpopulation of GAG-expressing VICs was observed to localize at the tip and hinge regions next to *Hapln1*^+^ VECs, which were located in the area where leaflet coaptation occurs^16^. Nevertheless, the study did not delve deeper into the cellular and molecular mechanisms driving ECM maturation and valve remodeling. Specifically, how different VEC subtypes that are exposed to differing flow regulate VIC physiology and ECM maturation, thereby influencing valve remodeling.

To determine the cellular composition of human fetal valves and elucidate the molecular mechanisms underlying human valve remodeling, we performed scRNA-seq to extensively profile all four human fetal valves during the early valve remodeling phase (gestational week 15). We identified a novel Elastin-producing *APOE^+^* VIC subtype, and revealed the function of these cells in regulating elastogenesis during early-stage valve remodeling via APOE-facilitated NOTCH activation. This study provided a high-resolution cell atlas of human fetal valves, and the opportunity to understand the pathogenesis of elastic malformation in various congenital valve diseases.

## Methods

### Human Heart Tissue Collection

De-identified human fetal heart tissues were obtained in collaboration with the University of Washington Birth Defects Research Lab (BDRL, IRB STUDY00000380), University of Southern California and Children’s Hospital Los Angeles, and Cincinnati Children’s Research Foundation (CCRF, IRB 2011-2856). All samples were collected from patients who provided signed informed consent according to IRB guidelines in accordance with all university, state, and federal regulations. The only clinical information collected was gestational age and the presence of any maternal or fetal diagnoses. Fetal tissues for scRNA-seq and primary cell culture were transferred from BDRL to Stanford University under Material Transfer Agreement #44556A. The samples, data, and/or services derived from CCRF were provided by Discover Together Biobank. The above studies were classified as exempt because of the deidentified nature of the valve samples. Clinicopathological baseline characteristics of patients were shown in Supplemental Table 1.

### Single-cell RNA-seq on Human Fetal Heart Valves

One normal human fetal heart (week 15, male) and one heart sample with Pulmonary Stenosis judged by ultrasound showing underdeveloped right heart and abnormal right outflow tract (week 15, male) were manually micro-dissected. Aortic, pulmonary, mitral, and tricuspid valves were collected, minced, and digested using the same digestion buffer as VECs and VICs isolation. Following digestion for 10 min at 37°C, EGM-2 medium was added to stop the reaction and cells were transferred through a 40 μm cell strainer. Cells with viability>90% were sent to Stanford Functional Genomics Facility (SFGF) for 10X chromium single-cell RNA-seq paired-end library preparation using V2 version (10X Genomics). All samples were uniquely indexed, mixed, and evenly distributed into the Illumina HiSeq 4000 for sequencing.

### Quantification and Statistical Analysis

All data represent mean ± SEM. Statistical significance was determined by parametric tests (all data possesses equal variance p>0.05): unpaired 2-tailed t-test (2 groups), ANOVA (>2 groups) with post hoc tests as indicated; or non-parametric test: Mann-Whitney (2 groups). A p value of <0.05 was considered significant. For all cell culture experiments, n represents different cell lines or combinations of EC and VIC from different patients or control subjects. For all image quantification, n represents different specimens from different patients or control subjects.

## Data Deposition and Availability

The raw and processed data from single-cell RNA sequencing in this study have been deposited with the Gene Expression Omnibus (https://www.ncbi.nlm.nih.gov/geo/) under accession number GSE228638.

All detailed description of methods used in this study are included in the Supplemental Materials.

## Results

### scRNA-seq Revealed VIC Subpopulations in Human Fetal Valves

To dissect the cellular and molecular mechanisms regulating the early events during valve remodeling, human fetal hearts from week 15 (W15) of gestation were examined. Movat staining of aortic, pulmonary, mitral, and tricuspid valves revealed enriched GAG (blue) compared with collagen (yellow) or elastin (black), indicating the lack of well-defined tri-laminar structures typically observed in mature valves and confirming that valvular remodeling was still occurring at this developmental stage (**Figure 1A**). Through microdissection of the W15 human fetal heart, four valves were collected for scRNA-seq profiling. A total of 35,081 cells were used for downstream analysis. After Quality Control and employing CCA algorithm to eliminate batch effect^17^ (**Figure S1A-B**), six major cell types were identified based on a group of well-established cell-type specific markers being used within previous annotation in rodent valves^16^ – VICs (*COL1A1,* cluster 1), VECs (*PECAM1,* cluster 2), erythroid cells (*HBA1,* cluster 3), neural-related cells (*SOX10,* cluster 4), macrophages (*CD74,* cluster 5), and dendritic cells (*KLRB1,* cluster 6) (**Figure 1B**, **S1C-D**). The heterogeneity of VICs and VECs in cardiac valves have previously been reported^16^. Given the significant roles of VICs in regulating valvular remodeling^1, 18^, we further subclustered the VIC population and revealed five subtypes: Elastin-VICs (*ELN*^+^, *APOE*^+^), CLDN11-VICs (*CLDN11*^+^), Collagen-VICs (*FMOD*^+^), Glycosaminoglycan-VICs (GAG, *LUM*^+^), and proliferative (*MKI67*^+^) VICs, each with their unique transcriptional signatures (**Figure 1C–D**, **S1E-F**).

**Figure 1:**
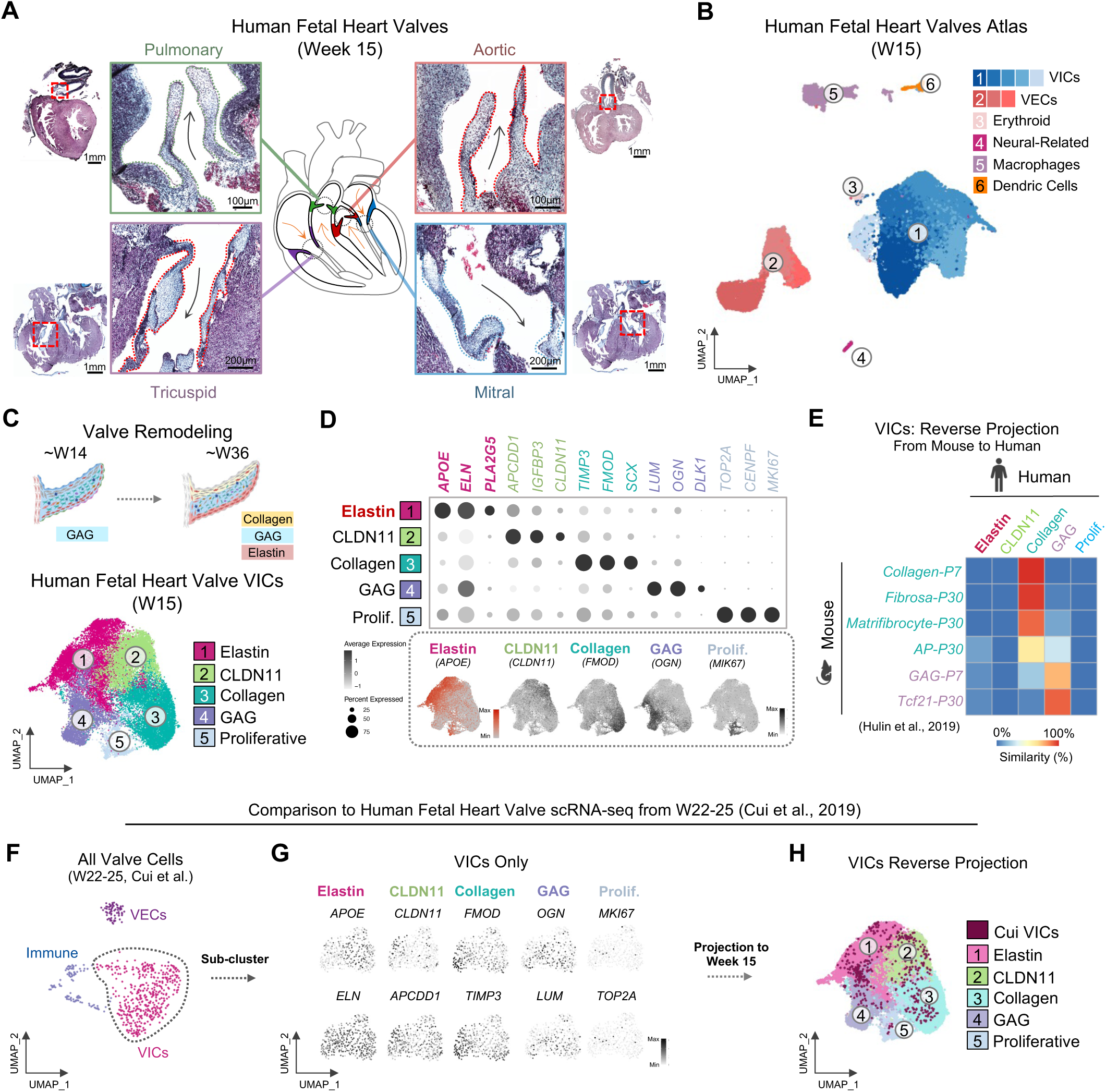
scRNA-seq Discovered Novel Elastin-Expressing VICs In Human Fetal Heart Valves. A. Representative Pentachrome staining of four cardiac valve structures from a week 15 normal human fetal heart. Pentachrome stain displays mostly glycosaminoglycan ECM (GAG) (blue). Elastin: black. Collagen: yellow. Muscle: red; **B.** Uniform Manifold Approximation and Projection (UMAP) visualization of 35081 cells isolated from four different valves of one W15 healthy human fetal heart; **C.** UMAP visualization of five VIC subtypes; **D.** Dot plot (upper) and feature plots (lower) for marker genes within each VIC subtype; **E.** Heatmap showing the similarity of each VIC subtype between human and mouse through reverse projection; **F.** UMAP visualization of valve cell types from W22-25 healthy human fetal valves (Cui et al., 2019); **G.** Feature plots of defined VIC marker genes within W22-25 VICs (Cui et al., 2019); **H.** UMAP demonstrating the reverse projection of VICs from W22-25 (Cui et al., 2019) to W15. VICs: Valve Interstitial Cells; VECs: Valve Endothelial Cells; GAG: Glycosaminoglycan; Prolif.: Proliferative; AP: Antigen-Presenting; P7: postnatal day 7; P30: postnatal day 30.

To understand the differences between human and mouse valves, we compared our human VIC dataset to a publicly available postnatal mouse valve scRNA-seq dataset with well-defined VIC subtypes^16^. Reverse projection was performed to calculate the similarity of expression profiles for each VIC subtype. The human GAG-VICs exhibited similar expression patterns as mouse postnatal day 7 (P7) GAG-VICs and P30 Tcf21^+^ VICs, while the human Collagen-VICs shared a similar transcriptomic profile with mouse P7 Collagen-VICs, P30 Fibrosa, Matrifibrocyte, and Antigen Presenting (AP)-VICs. Intriguingly, the reverse projection analysis revealed three novel VIC subtypes during the dynamic valve remodeling phase that were not described in rodent valves-Elastic-VICs, CLDN11-VICs and Proliferative VICs (**Figure 1E**, **S2A-E**). The Proliferative VICs were enriched for *MKI67*, indicating a relatively early stage of valve remodeling^2^. CLDN11-VICs were enriched for a tight junction gene *CLDN11* (*Claudin 11*)^19^ and a known Wnt inhibitor *APCDD1* (*APC down-regulated 1*)^20^. Elastin-VICs were highly enriched for *ELN* and two lipoprotein-related genes: *Phospholipase A2 Group V* (*PLA2G5*), a calcium-dependent phospholipase, and *Apolipoprotein E (APOE)*, an apolipoprotein that serves as a carrier of lipids and cholesterol^21, 22^. However, the role of APOE in valve remodeling is completely unknown. In this study, we first focused on the subset of Elastin-producing VICs, as elastin fiber formation is one of the key events during valve remodeling, and their transcriptomic signatures have yet to be elucidated at single cell resolution.

To validate the presence of Elastin-VICs across different stages of human valve development, we compared our data with an existing human fetal valve dataset generated during late valve remodeling phase (Cui et al., week 22-25)^14^. Elastin-VICs were also present in Cui’s VICs dataset^14^ based on the marker genes we have identified above (*ELN, APOE*, and *PLA2G5*) (**Figure 1F–G**, **S2F-H**), although their VICs population barely clustered into similar distinct subgroups, possibly due to the low cell number input (**Figure 1G, Table 1**). Through reverse projection, we confirmed that the Elastin-VICs from W22-25 valves sparsely scattered on W15 valve UMAP (**Figure 1H**), suggesting the presence of Elastin-VICs during various stages of human valve remodeling.

**Table 1:** Cell Numbers Captured in Each scRNA-seq Dataset Used in the Current Study

| Dataset | Source | Cell Type |  |  |  |  |  | Total |
| --- | --- | --- | --- | --- | --- | --- | --- | --- |
|  |  | VICs | VECs | Immune |  | Neural Related | Erythroid |  |
|  |  |  |  | Macrophages | Dendric Cells |  |  |  |
| Human (W15) | Current | 25853 | 6449 | 2140 | 348 | 173 | 118 | 35081 |
| Mouse (P7+P30) | Hulin et al. | 3239 | 459 | 412 |  | 27 | 0 | 4137 |
| Human (W22-25) | Cui et al. | 430 | 72 | 106 |  | 0 | 0 | 608 |

### APOE is Essential for Elastogenesis During Valve Remodeling

*APOE* was found to be the top enriched gene in the Elastin-VICs population. While numerous studies have linked *APOE* to valvular calcification in senile valves^23–25^, the role of APOE in VIC physiology during fetal valve elastogenesis has not been studied. Given the co-expression of *APOE* and *ELN* in Elastin-VICs, we first examined the role of APOE in regulating elastogenesis, which is critical for valvular remodeling, and maturation. Through RNA *in situ* hybridization and immunostaining, we validated the expression of *APOE, ELN,* and additional elastogenesis-related genes such as *EMILIN1*^3^ in four human fetal valves collected at the early valve remodeling stage (W15-17). Interestingly, we found that these Elastin-VICs were specifically located underneath VECs in the direction of unidirectional flow (**Figure 2A–B**, **S3A**), which is consistent with previous findings regarding the location of early, non-cross-linked elastin in human valves^3^. To further study the role of *APOE* in regulating the function of Elastin-VICs, particularly in elastogenesis (**Figure 2C**), we isolated primary valvular cells from human fetal hearts. Compared with VECs, VICs showed high levels of COL1A1, αSMA, and low levels of endothelial marker PECAM1, judged by qPCR, immunostaining, and flow cytometry (**Figure S3B-D**). *APOE* expression was then perturbed in primary human fetal VICs using small interfering RNAs (siRNAs) (**Figure S3E**). Significant downregulation of genes associated with elastogenesis such as *FBN1*, *LTBP1*, *EMILIN1*, and *LOXL1* was observed in APOE-deficient VICs although the expression of *ELN* remained unchanged (**Figure 2D**, **Figure S3F**). Protein level analysis revealed significant decrease of EMILIN1 in APOE-deficient VICs by immunostaining (**Figure 2E**). Notably, during fetal valve remodeling stage, an αSMA^+^ activated myofibroblasts underneath endocardium below the outflow surface were previously reported to be engaged in matrix remodeling^1^. Due to the similarity of spatial location between Elastin-VICs and the activated myofibroblast, and an *ACTA2* enrichment in Elastin-VICs (**Figure S3G**), we further tested whether Elastin-VICs might gain a “myofibroblast” phenotype. By performing *in vitro* contraction assay using collagen gel, we found that APOE-deficient VICs exhibited impaired contraction compared with the scramble group (**Figure S3H**). This observation was accompanied by a significant decrease in expression of contraction-related genes (*ACTA2*, *MYH11*)^26, 27^ (**Figure S3I**), all suggested a gain of myofibroblast phenotype within Elastin-VICs. In summary, APOE^+^ Elastin-VICs were spatially located underneath VECs sensing unidirectional flow and played important roles in regulating elastogenesis during valve remodeling.

**Figure 2:**
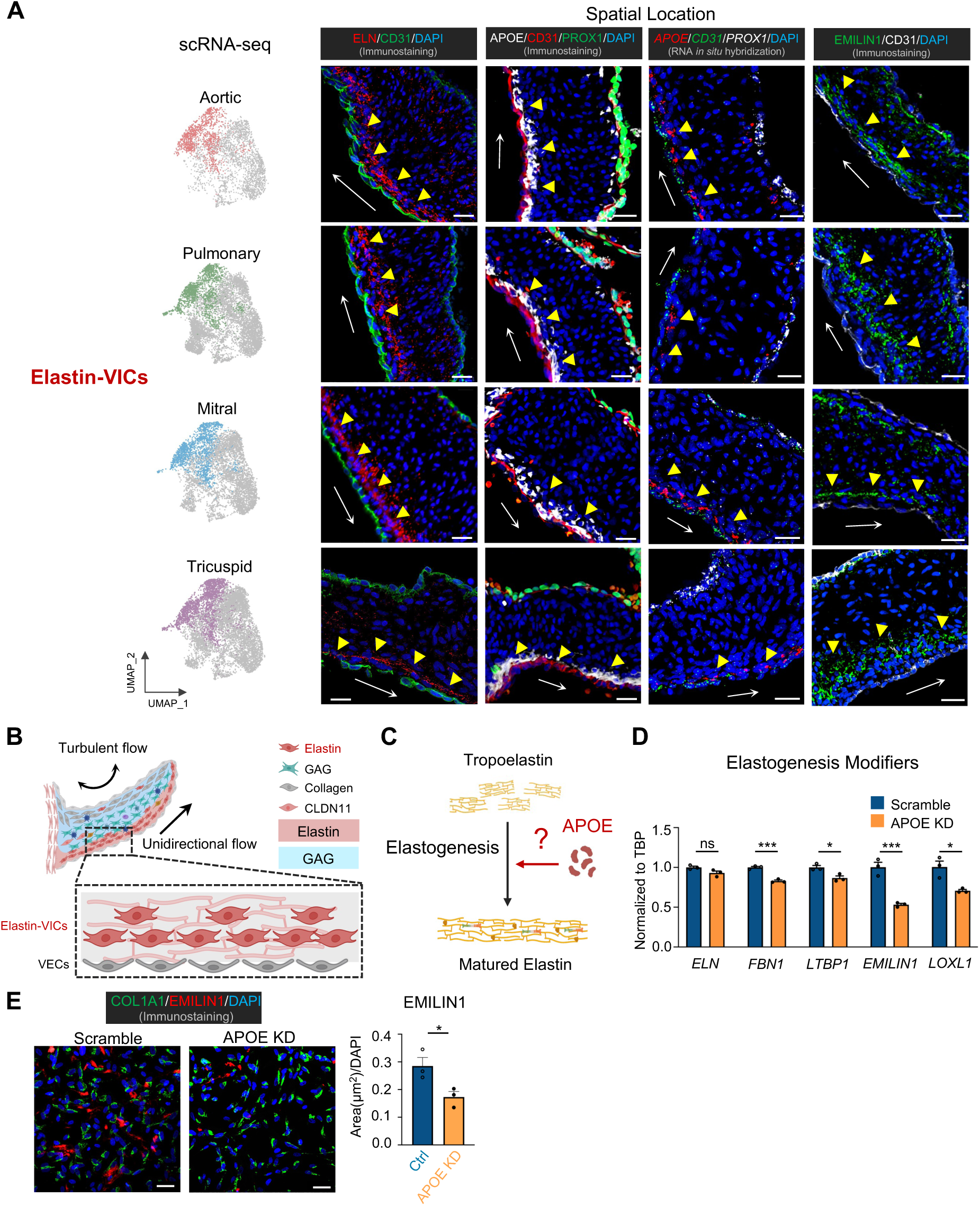
APOE Is Essential For Elastogenesis in Elastin-VICs. A. Left: UMAP distribution of Elastin-VICs within each valve. Right: Representative immunofluorescence staining of Elastin, APOE, EMILIN1, and RNA *in situ* hybridization of *APOE*, in age-matched four human fetal valves (age from W15-17). n=3. Scale bar: 20μm. Arrows represent unidirectional flow directions; **B-C.** Illustration figures highlighting spatial location of Elastin-VICs within human fetal heart valve (**B**) and demonstrating potential role of APOE in the elastogenesis of VICs (**C**); **D.** qPCR analysis of elastogenesis related genes expression in cultured human VICs. n=3 biological repeats. Data shown as the mean ± SEM. ns p>0.05, *p<0.05, ***p<0.001, scramble vs APOE KD; **E.** Left: Representative immunofluorescence staining of EMILIN1 and COL1A1 in cultured human VICs. Scale bar: 50μm; Right: Quantification of positive staining area of EMILIN1 per cell in VICs. n=3; Statistics in D: Unpaired 2-tailed t-test (2 groups). KD: Knockdown.

### APOE-deficiency and Elastin Fragmentation in Pulmonary Valves with Pulmonary Stenosis

Elastin fragmentation is a common pathological feature of myxomatous valves^28^, which appeared to be one of the most common pathological presentations seen in congenital valve diseases such as mitral regurgitation, bicuspid aortic valve and Tetralogy of Fallot (TOF) with PS^29–32^. To further expound on the role of *APOE* in the pathogenesis of congenital valve diseases, we compared normal pulmonary valves (PVs) versus age-matched PVs exhibiting Pulmonary Stenosis (PS), a congenital heart disease characterized by abnormal valve formation through scRNA-seq and histology staining (**Figure 3A**). Significant abnormality in ECM composition was revealed in PS valves through Pentachrome staining, which showed an accumulation of GAGs and a decrease in elastin layer thickness compared to age-matched control valves (**Figure 3B**, **Table S1**). By performing immunofluorescence on elastin in both control and PS valves, severe loss of *ELN* expression, coupled with disrupted elastin fiber structures were observed in PS valves when compared to age-matched controls. This suggests a profound elastogenesis defect within the PS valves. Intriguingly, a drastic reduction of APOE, as well as EMILIN1, LOXL1, and αSMA were observed in PS valves at protein levels (**Figure 3B–C**, **S4A**), suggesting a pathogenic association of APOE in PS. scRNA-seq comparing control and age-matched PS PVs was then performed to comprehensively understand the role of APOE and Elastin-VICs in regulating elastogenesis and valve remodeling. After annotating major valvular cell types (**Figure S4B-C**), and VIC subtypes (**Figure 3D**, **Figure S4D**), the decreased expression of *APOE* and *ELN* in Elastin-VICs were observed (**Figure 3E**); this was accompanied by the reduction of elastogenesis-related genes including *FBN1, LTBP1, EMILIN1,* and *LOXL1*. In addition, contractile related genes *ACTA2*, *MYH11*, and *CNN1* were significantly downregulated in PS Elastin-VICs (**Figure 3F**), similar to what was observed when *APOE* was knocked down in fetal VICs (**Figure 2D**, **Figure S3I**). Collectively, we found that PS PVs displayed distinct elastin fragmentation and reduced expression of *APOE* in Elastin-VICs, suggesting a critical role of APOE-regulated elastogenesis defect in human PS pathology.

**Figure 3:**
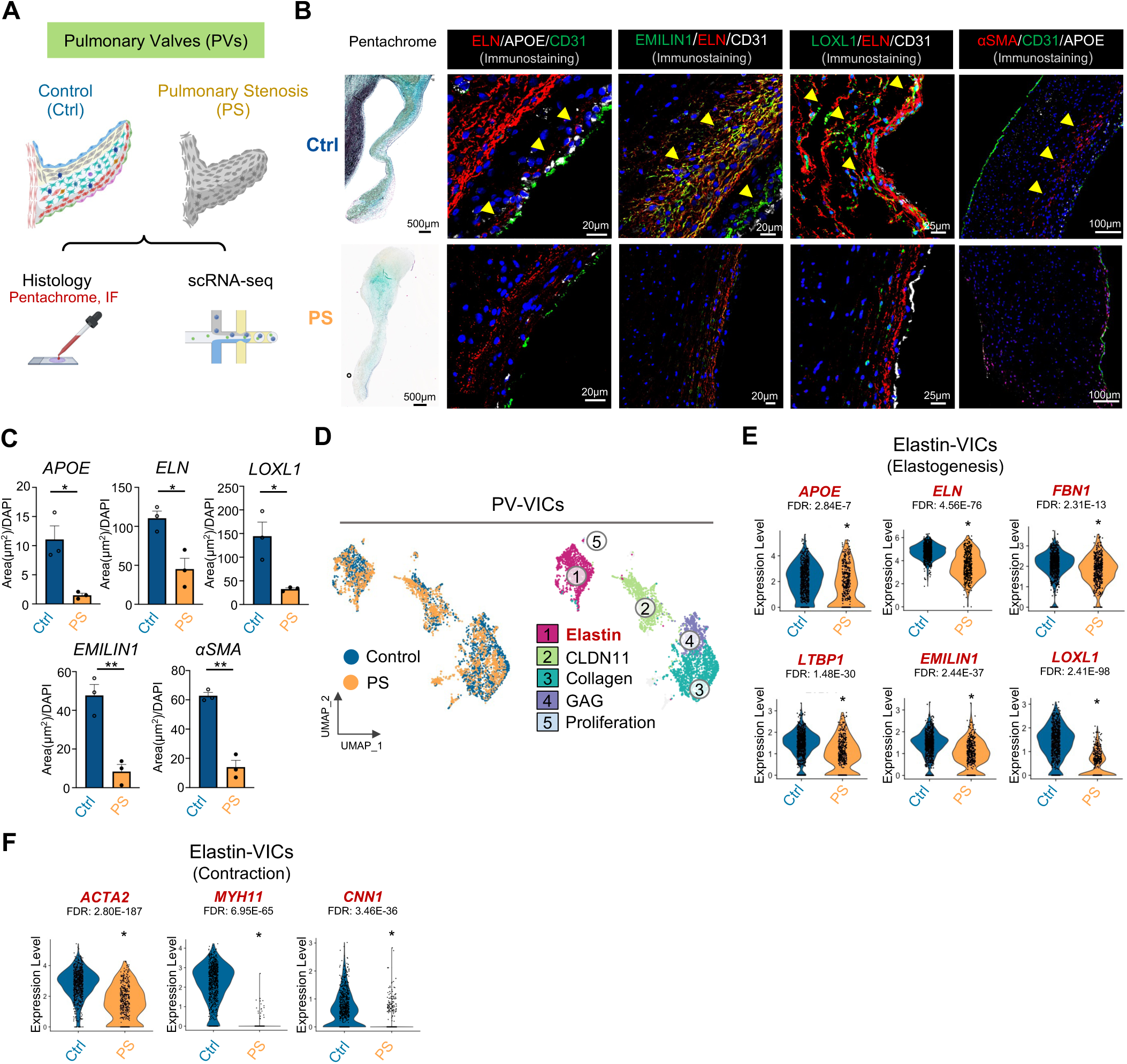
APOE-deficiency and Elastin Fragmentation in Pulmonary Valves with Pulmonary Stenosis. A. Methods used to explore the differences between postnatal control pulmonary valves (PVs) and pulmonary stenotic (PS) valves; **B.** Pentachrome (Left) and immunofluorescence staining (Right) of ELN, APOE, EMILIN1, LOXL1 and αSMA in PVs from control and PS. Age from 5 months to 10 years. n=3 (Table S1); **C.** Quantification of positive staining area of APOE, ELN, LOXL1, EMILIN1 and αSMA per cell within elastin layer of PV between healthy controls and PS. n=3; **D.** UMAP of PV-VICs from control vs. patient with PS; **E-F.** Violin plots of *APOE*, elastogenesis (**E**), and contraction (**F**) related genes expression within PV Elastin-VICs comparing control and PS. Data are shown as the mean ± SEM. *p < 0.05, **p < 0.01. Statistics in C: Unpaired 2-tailed t-test. IF: Immunofluorescence, Ctrl: Control, PS: Pulmonary Stenosis, PVs: Pulmonary Valves, FDR: False Discovery Rate.

### NOTCH Ligands Secreted by Unidirectional VECs Regulates Elastogenesis in Elastin-VICs

VICs are spatially juxtaposed to VECs in valve leaflets, however cellular interactions between VICs and VECs remain to be discerned. It has been shown that loss of VEC-specific factors stifled VICs growth, therefore resulting in valve defects^9^. Since Elastin-VICs spatially located adjacent to VECs under unidirectional flow, we postulate that in addition to APOE, external factors originating from VECs could also play a part in regulating elastogenesis in Elastin-VICs. The VEC populations were first extensively profiled in all four human fetal valves. Three VEC subtypes with distinct transcriptomic signatures were identified – Lymphatic VECs (*PROX1^+^, FOXC2^+^*), CD55-VECs (*DKK2^+^, CD55^+^*), and *prostaglandin D2 synthase* (*PTGDS*) expressing PTGDS-VECs (*PTGDS^+^, KRT18^+^*) (**Figure 4A–B and Figure S5A-C**). Notably, the PTGDS-VEC subtype has not been described in previous mouse VEC dataset^16^ (**Figure 4C and Figure S5D-G**). Immunofluorescence staining and RNA *in situ* hybridization confirmed the presence of CD55-VECs and PTGDS-VECs in all four human fetal valves, and spatially located under unidirectional flow (**Figure 4D**, **Figure S5J-K**). Conversely, FOXC2^+^ Lymphatic VECs were located adjacent to turbulent flow (**Figure S5I-J**). Therefore, both CD55-VECs and PTGDS-VECs were coined unidirectional VECs (UDVECs), spatially juxtaposing the Elastin-VICs (**Figure S6A**). Of note, higher proportions of PTGDS*-*VECs were found in semilunar (aortic and pulmonary) valves compared to atrioventricular valves (mitral and tricuspid), while Lymphatic VECs were found higher in proportion in atrioventricular valves (**Figure 4D**, **and Figure S5B, H, K**), indicating different cellular compositions in semilunar valves versus atrioventricular valves, possibly due to different flow patterns^14^.

**Figure 4:**
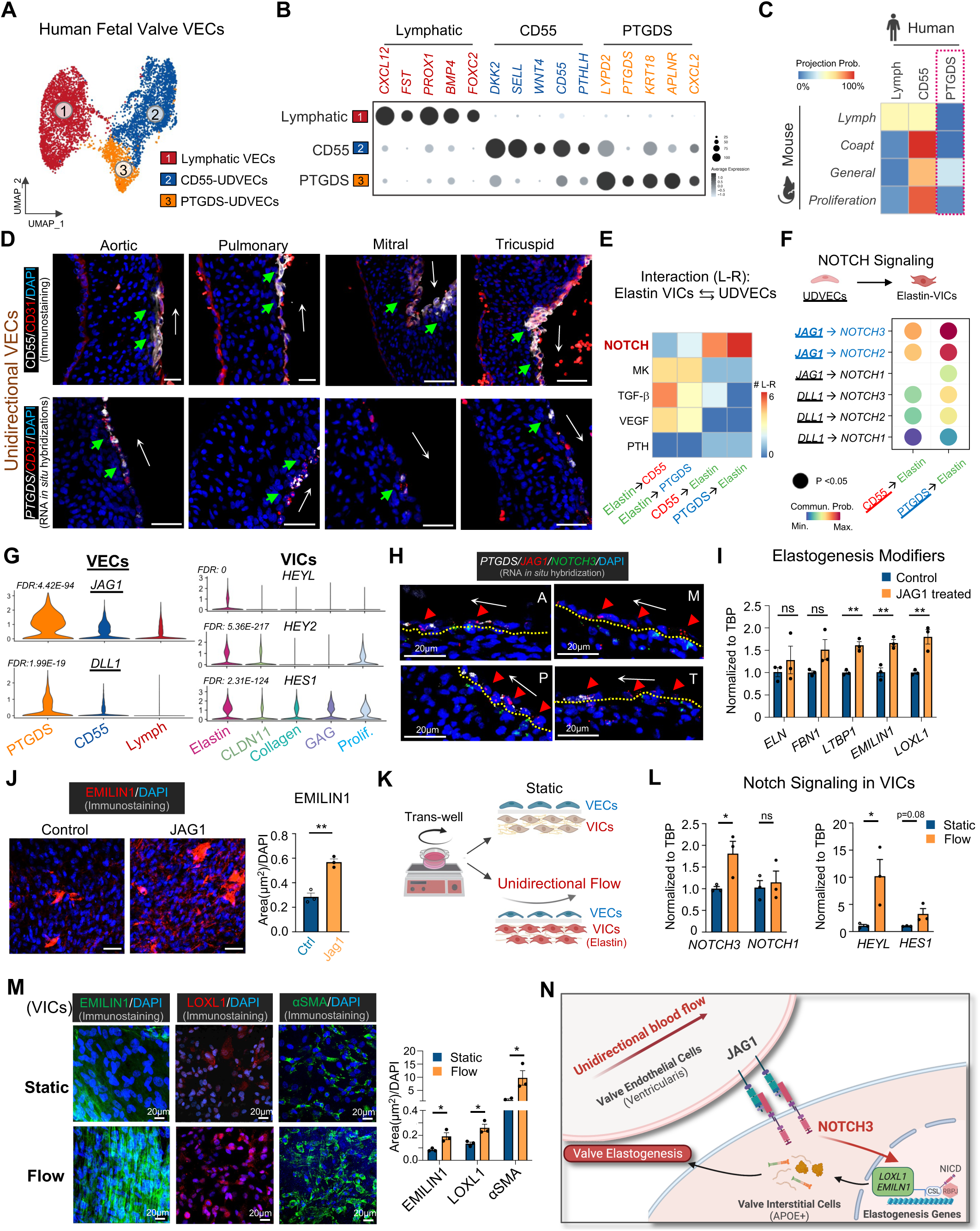
Notch Ligands from Unidirectional VECs Regulates Elastogenesis in Elastin-VICs. A. UMAP visualization of VEC subtypes in four control valves; **B.** Dot plot for marker genes within each VEC subtype; **C**. VEC subtypes similarities between human and mouse through reverse projection; **D.** Immunofluorescence staining of CD55 and RNA *in situ* hybridization staining of *PTGDS* in four human fetal valves at W15. n=3, Scale bar: 20μm. White arrows represent unidirectional flow directions; **E.** Top non-ECM related Ligand-Receptor pairs between Elastin-VICs and unidirectional flow-sensing VEC subtypes; **F.** Major Notch Ligand-Receptor pairs captured from UDVECs to Elastin-VICs; **G.** Violin plots of Notch-related ligands and downstream targets in VEC and VIC subtypes. FDR: PTGDS-VECs vs. other VEC subtypes or Elastin-VICs vs. other VIC subtypes; **H.** RNA *in situ* hybridization of *JAG1* and *NOTCH3* in four human fetal valves, white arrow represents unidirectional flow directions, red arrowhead indicated *JAG1*^+^ VECs juxtaposed to *NOTCH3*^+^ VICs; **I.** qPCR analysis of elastogenesis related genes in VICs with Jag1 treatment; **J.** Left: Representative immunofluorescence staining of EMILIN1 in cultured human VICs. Scale bar: 50μm; Right: Quantification of positive staining area of EMILIN1 per cell in VICs; **K.** Demonstration figure showing co-culture of VECs and VICs under unidirectional or static flow conditions; **L.** qPCR analysis of Notch signaling related genes in VICs co-cultured with VECs under static or unidirectional flow; **M.** Immunostaining of EMILIN1, LOXL1 and αSMA in VICs co-culture with VECs under static or unidirectional flow. Unidirectional shear stress: 8.52 dynes/cm^2^. Jag1: 15ug/ml. **N.** Demonstration of unidirectional flow-induced Jag1 production by VECs, which activates Notch and promote elastogenesis in VICs. n=3 biological repeats in each group. Data shown as the mean ± SEM. ns p>0.05, *p<0.05, **p<0.01. Statistics in I, J, L, M: Unpaired 2-tailed t-test (2 groups). UDVECs: Unidirectional VECs, ECM: Extracellular Matrix, Lymph: Lymphatic.

Next, CellChat analysis^33^ was employed to determine and characterize cell-cell interactions between spatially-adjacent VICs and VECs. As expected, ECM-related signaling pathways such as collagen and laminin were found to be significantly enriched in mediating the crosstalk between Elastin-VICs (source) to UDVECs (receivers) (**Figure S6B**). This further confirms the involvement of Elastin-VICs in supporting VECs through ECM-related interactions^34^. Interestingly, our data also demonstrated that UDVECs provided ligands to active key signaling pathways in Elastin-VICs. In total, 158 ligand-receptor (L-R) pairs were identified between CD55-VECs (source) and Elastin-VICs (receiver), and 137 pairs were identified between PTGDS-VECs (source) and Elastin-VICs (receiver) (**Figure S6C**). Among all the non-ECM-related pathways, Notch signaling was found to be the most enriched pathway present between the UDVECs (source) and Elastin-VICs (receiver) (**Figure 4E**, **Figure S6D, Figure S7**). Specifically, Notch ligands *DLL1* and *JAG1* were found to be highly expressed by UDVECs compared to Lymphatic VECs under the turbulent flow, which activated Notch downstream genes in Elastin-VICs (**Figure 4F–G**). In particular, the *JAG1-NOTCH3* L-R pair was the most prevalent (**Figure 4F**, **Figure S6G**). We also noted that between both UDVEC subtypes, PTGDS-VECs expressed relatively higher levels of *JAG1* and *DLL1*, while *NOTCH1* and *NOTCH3* were enriched in Elastin-VICs (**Figure 4G**, **Figure S6G**). Consistent with this observation, gene ontology (GO) analysis showed enrichment of NOTCH signaling in PTGDS-UDVECs and Elastin-VICs (**Figure S6E-F**). RNA *in situ* hybridization in human valve tissues further validated that *JAG1* was enriched within UDVECs while *NOTCH3*-expressing cells were found juxtaposed to these *JAG1^+^* cells (**Figure 4H**). This indicates that UDVECs might be central in activating Notch signaling in Elastin-VICs.

To further investigate the role of *JAG1* in Elastin-VICs physiology and function, human fetal VICs were treated with exogenous Jag1 *in vitro*. We proved that exogenous Jag1 activated Notch signaling in VICs evidenced by increased *NOTCH3*, *HEYL* and *HEY2* expression (**Figure S6H**). More importantly, Jag1 treatment drastically induced the expression of elastogenesis-related genes such as *FBN1*, *LTBP1*, *EMILIN1*, and *LOXL1,* although the expression of *ELN* remained unchanged (**Figure 4I**, **Figure S6I**). Protein level analysis revealed that Jag1 treatment significantly increased EMILIN1 expression (**Figure 4J**), further supporting the role of Jag1 in promoting elastin assembly. Notably, Jag1 treatment also enhanced contractility of fetal VICs (**Figure S6J**) and induced the upregulation of contractility-related genes *ACTA2*, *MYH11*, and *TAGLN* (**Figure S6K**). These data further underscore the function of Jag1 in promoting the activation of myofibroblast phenotype within Elastin-VICs.

Since Elastin-VICs were spatially adjacent to JAG1-expressing UDVECs, which were exposed to unidirectional flow, we hypothesized that unidirectional flow was responsible for JAG1 induction in UDVECs. To test this hypothesis, we first isolated human fetal VECs, characterized by high expression levels of *PECAM1* and *CDH5*, and low levels of VIC markers COL1A1 and αSMA (**Figure S6L-N**). In the presence of unidirectional flow, an increase in *JAG1* coupled with elevated expression of flow-responsive genes such as *KLF2*, *KLF4* and *NOS3* were observed when compared to VECs cultured under static conditions (**Figure S6O**). To further understand the Notch-mediated VEC-VIC crosstalk in valve elastogenesis, fetal VECs were co-cultured with VICs in the presence of unidirectional flow (**Figure 4K**). Notch activation was observed in VICs as evidenced by the presence of Notch signaling target genes (**Figure 4L**). Similar to Jag1-treated VICs, expression levels of elastogenesis-related proteins such as EMILIN1 and LOXL1 were significantly elevated in VICs cultured under the unidirectional flow compared to the static condition. Additionally, αSMA expression increased significantly, suggesting an activated myofibroblast phenotype in VICs underneath unidirectional flow^27^ (**Figure 4M**). Altogether, we found that unidirectional flow induced *JAG1* expression in UDVECs, which signaled through the NOTCH3 receptor in neighboring Elastin-VICs, thereby activating Notch signaling in Elastin-VICs. This in turn promoted elastogenesis in Elastin-VICs (**Figure 4N**).

### Impairment of Notch signaling in Pulmonary Valves with Pulmonary Stenosis

To further understand the role of JAG1-NOTCH3 signaling in the context of congenital valve disorders, we examined the expression levels of Notch ligands and receptor in PS VECs and VICs. We found that *JAG1* expression was significantly downregulated in PTGDS-VECs and CD55-VECs of PS PVs compared with healthy controls, suggesting that *JAG1* insufficiency in UDVECs might contribute to elastogenesis defect in PS (**Figure 5A–B**, **Figure S8A-C**). Next, GO enrichment analysis of genes down-regulated in PS Elastin-VICs revealed that a large proportion of those genes were involved in the Notch signaling cascade (**Figure 5C**); the expression of the Notch receptor *NOTCH3* and downstream target *HEYL, HES1 and JAG1* were significantly reduced in PS Elastin-VICs compared with healthy control (**Figure 5D**, **Figure S8D**). This was further confirmed by the significant decreased expression of *JAG1* and *NOTCH3* in three additional PS valves (**Figure 5E**). These data reinforced the importance of Notch-mediated VEC-VIC crosstalk in facilitating normal valve development.

**Figure 5:**
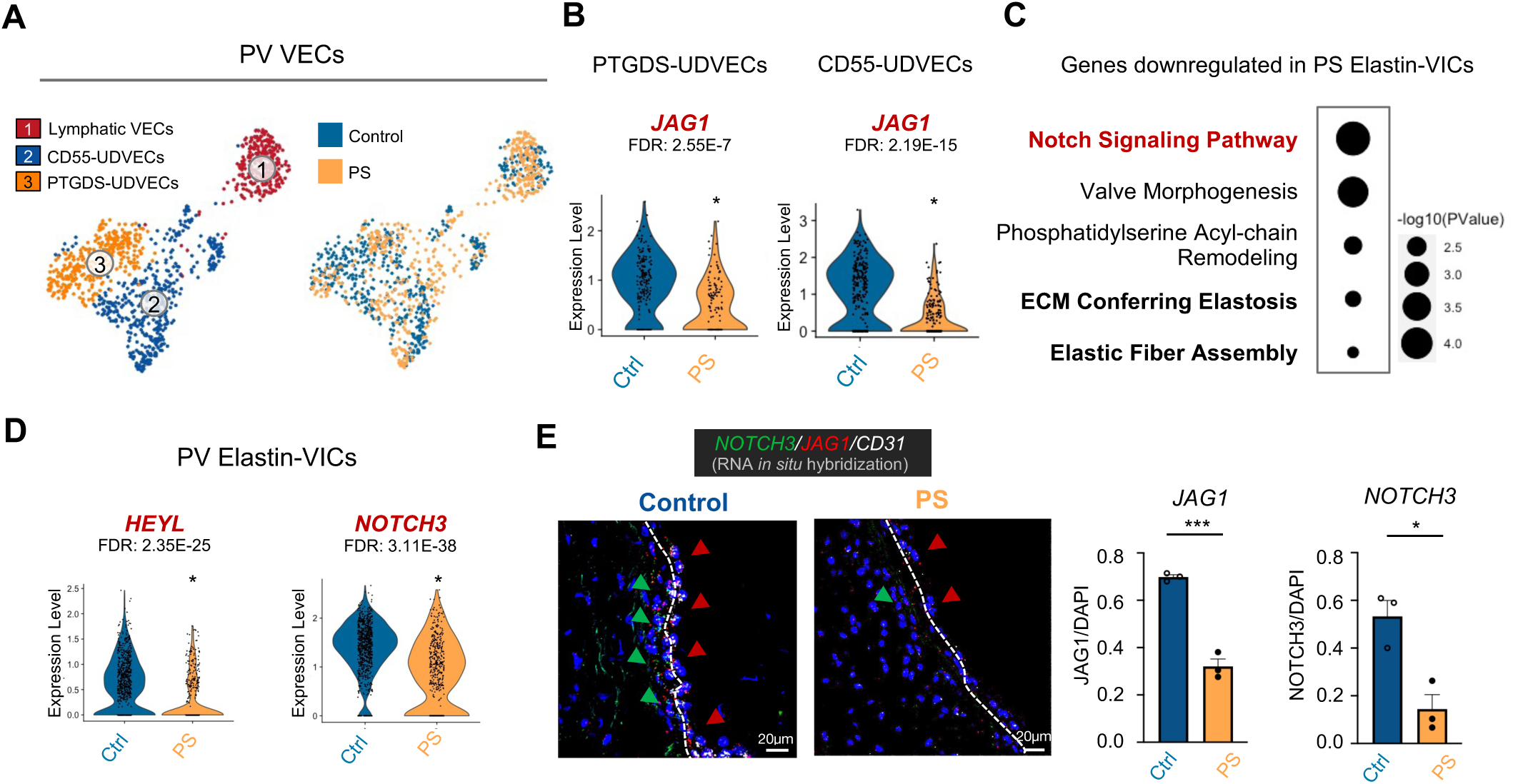
Impairment of Notch Signaling in Pulmonary Valves with Pulmonary Stenosis. A. UMAP visualization of cell subtypes (Left) and cell distribution (Right) within control and pulmonary stenosis (PS) VECs; **B.** Violin plots comparing *JAG1* expression between control vs. PS in *PTGDS-*VECs and *CD55-*VECs; **C.** GO analysis of genes downregulated in PS Elastin-VICs; **D.** Violin plots comparing *HEYL* and *NOTCH3* expression between control vs PS in Elastin*-*VICs; **E.** Left: RNA *in situ* hybridization of *JAG1* (red arrow) and *NOTCH3* (green arrow) in pulmonary valves (PV) from control vs. PS. Right: Percentage of *JAG1^+^* cells in UDVECs and *NOTCH3^+^* VICs in Elastin layer VICs in valve tissues from control vs. PS. n=3 different samples in each group. Data shown as the mean ± SEM. *p<0.05, **p<0.01, ***p<0.001. Statistics in E: Unpaired 2-tailed t-test (2 groups).

### APOE is Indispensable for Notch activation in Elastin VICs

Hitherto, we have shown that APOE expression and Notch activation in human fetal Elastin-VICs is critical for valve elastogenesis. However, the relationship between APOE and Notch signaling in VICs remains ambiguous; it is unclear whether the pro-elastogenic effects of both mediators were interdependent or independent (**Figure 6A**). To elucidate this, we first investigate the potential interplay between APOE and Notch signaling. We found that APOE knockdown in VICs dampened the activation of Notch signaling to a similar level as Gamma Secretase Inhibitor (GSI) treatment (**Figure 6B**, **S9A**), suggesting that APOE is required for Notch activation in VICs. Additionally, VICs with APOE suppression as well as Notch inhibition resulted in impaired activation effect of Jag1 in the expression of elastogenesis-related genes (**Figure 6C**). Protein level analysis revealed that Jag1 treatment in VICs with APOE knockdown decreased tropo-elastin and elastin amount (**Figure 6D**), and impaired the effect of Jag1 in increasing EMILIN1 expression (**Figure 6E**). Notably, impairment of the effect of Jag1 in the enhancement of contractility of VICs (**Figure S9B**), accompanied by reduced expression of contractile genes were also discovered (**Figure S9C**) by treating Jag1 to VICs with APOE knockdown, suggesting that APOE is critical in maintaining Jag1 induced myofibroblast phenotype. All together, these data suggested that in VICs, APOE is indispensable for Notch activation, reinforcing the pivotal role of APOE-regulated Notch signaling in mediating elastogenesis in VICs (**Figure 6F**).

**Figure 6.**
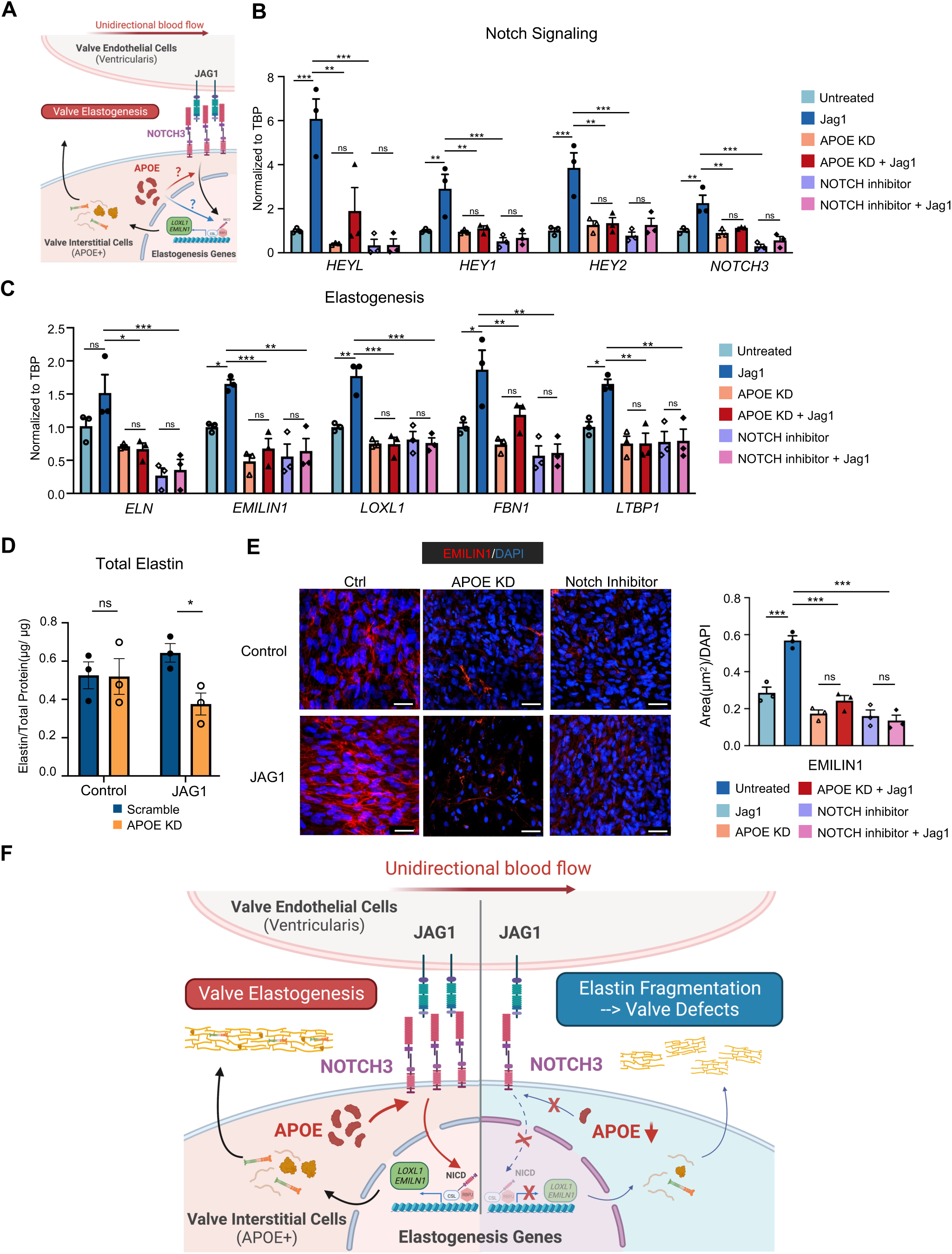
APOE is Required for Notch Signaling Activation. A. Illustration of interaction between APOE and Notch signaling in promoting elastogenesis; **B-C**, VICs were treated with different conditions. Jag1: 15ug/ml, L-685,458 (Notch inhibitor): 10uM; VICs within each treatment condition were subjected to qPCR analysis and immunostaining; **B.** qPCR analysis of Notch signaling pathway related genes in VICs; **C.** qPCR analysis of elastogenesis related genes in VICs; **D.** Quantification of total elastin (both tropo-elastin(soluble) and mature elastin (insoluble)) in VICs with APOE KD and/or Jag1 treatment. The elastin content was normalized to the total protein content in each condition; Data shown as the mean ± SEM. ns p> 0.05,*p<0.05,**p<0.01,***P<0.001. **E.** EMILIN1 immunostaining of VICs. Scale bar=50μm; **F.** Demonstration figure: On one side, unidirectional flow induced Jag1 production in VECs, which activated NOTCH3-mediated Notch signaling in Elastin-VICs and promote downstream elastogenesis-related genes expression. On the other side, APOE facilitated Notch signaling activation and elastogenesis genes expression. Insufficient APOE found in PS patients suppressed Notch signaling, which could not be rescued by Jag1 supplement. n=3 biological repeats in each group. Statistics in B-C, E: one-way ANOVA followed by Tukey’s test, D: Unpaired 2-tailed t-test (2 groups).

## Discussion

Great strides have been made over the past decade to understand the development of heart valves. However, the cellular and molecular mechanisms driving early valve remodeling during valvulogenesis remains unclear. Recent efforts to map the transcriptomic and cellular landscape of mammalian valves at various developmental stages have identified the presence of diverse VIC and VEC cell types^14–16^, however, the role of these cells in regulating ECM organization and valvular remodeling has not been fully determined. In this study, we discovered novel VIC and VEC populations in four different human fetal valves, discerned the relevance of these cell populations to valve development and disease, and uncovered critical VIC-VEC interactions that are essential for valve remodeling.

Firstly, we established a high-resolution single cell atlas of all four normal human fetal heart valves. Due to a larger number of captured cells, we were able to perform more comprehensive analysis on each cell cluster than previously published human heart valve datasets. Notably, we revealed the presence of an elastin-producing VIC subtype in all four human valves. This novel subtype of VICs was not captured in rodent valves, likely due to technical limitations caused by the significantly smaller valves and high ECM content, and therefore low cell numbers, in mouse hearts. We observed that Elastin-VICs were spatially located in the ventricularis and atrialis layers sensing unidirectional flow and express high levels of APOE. APOE, a well-studied apolipoprotein, was recognized for its role in regulating lipid metabolism, and its mutation was one of the leading causes of type III hyperlipoproteinemia^35^. An aged APOE-deficient mouse model has been widely utilized to investigate aortic valve sclerosis and calcification^23, 24^. However, the role of APOE in valve remodeling and physiology during heart development has not been reported.

Secondly, we described a novel role of APOE in regulating elastogenesis in Elastin-VICs during early-phase remodeling. By perturbing APOE in primary fetal VICs, genes associated with elastogenesis such as linking proteins FBN1 and EMILIN, latent transforming growth factor (TGF) β binding proteins (LTBPs), and lysyl oxidase (LOX, LOXL1) were significantly downregulated *in vitro*. Collectively, these proteins were crucial for the aggregation and cross-linking of tropoelastin into mature elastin^36–38^. We also observed that APOE regulates ACTA2 expression, a marker of activated myofibroblasts presents during valve remodeling^1^. These findings support the role pro-elastogenic role of APOE during the early-phase prenatal valve remodeling process at week 15, during which elastin fibers were still non-cross-linked, and were not mature enough to be detected through Movat staining. Additionally, PLA2G5, another lipid-metabolizing enzyme, was observed to be specifically enriched in Elastin-VICs. PLA2G5 was reported to be a key regulator of elastin stability in aortic walls^39^. Altogether, although lipid metabolism was shown to be involved in the pathogenesis of valve sclerosis and calcification, lipid metabolism in Elastin-VICs appears to be essential in promoting elastogenesis during valve remodeling. It is plausible that specific lipid metabolites regulated by APOE or PLA2G5 such as cholesterol, oleic acid, and linoleic acid^39^ are directly involved in mediating elastin development during valvulogenesis, as oleic acid and linoleic acid were reported to maintain elastin stability in aorta^39^.

Additionally, the role of APOE in regulating Notch activation in Elastin-VICs has not been described previously. It is evident that lipids could affect the level and duration of Notch signaling by modulating Notch ligands and NICD mobilization^40^. Since APOE and PLA2G5 are lipid metabolism-related genes enriched in Elastin-VICs, it is possible that they indirectly control Notch activation in VICs through altering cholesterol lipid mobilization. The regulation of Notch-related genes may be influenced by APOE through epigenetic machinery, as it has been shown to undergo nuclear translocation to act as, either a transcription factor by binding to approximately 1700 gene promoter regions^41^, or regulate gene expression through histone modifications and DNA methylation^42, 43^. APOE3 and APOE4 have also been reported to interact with the histone deacetylase (HDAC) complex to regulate gene expression^43^, while HDACs positively regulated Notch signal transduction by controlling NICD acetylation levels and directly impacting NICD1 protein stability^44^. Therefore, APOE may modulate Notch activation through various mechanisms including lipid metabolism and epigenetic regulation. Additional work is required to discern the exact underlying machinery.

We also discovered a novel CLDN11^+^ VIC subpopulation (CLDN11-VIC) that were not captured in Hulin et al.’s mouse dataset. While the spatial location and function of CLDN11-VICs within human valves remain unknown, we note that they were enriched for genes such as *APCDD1, IGFBP3,* and *ADAMTS19,* which were associated with valvular structure and Wnt signaling pathway^9, 20, 45, 46^. The shear-stress sensing Wnt signaling pathway was known to play an important role in valve endocardial cushion remodeling by regulating mesenchymal cell proliferation and condensation^9^. Gain-of-function of Wnt significantly impaired the earlier phase maturation of heart valves^47^.

Therefore, we speculated that CLDN11-VICs might involve in valve remodeling by responding to Wnt signaling induced by shear-stress. Aside from VICs, VECs play an equally important role during valve development and remodeling. This was exemplified by perturbed endothelial to mesenchymal transition (Endo-MT) caused by dysfunctional ECs, resulting in valve malformations observed in hypoplastic left heart syndrome (HLHS)^48–50^. In our study, we found that Notch signaling between UDVECs and Elastin-VICs facilitates elastogenesis during valve remodeling through the JAG1-NOTCH3 L-R pair. VECs serve as the major responder to hemodynamic forces during valve remodeling^9, 51^, and unidirectional flow has been shown to upregulate JAG1 and Notch signaling both *in vitro* and *in vivo*^52–54^. JAG1 mutation has been linked to Alagille syndrome, with pulmonary valve stenosis in cases of Tetralogy of Fallot^55^. Similarly, EC-specific deletion of JAG1 in mice leads to valvular defects such as pulmonary stenosis^56^. Previous studies also showed that the NOTCH3 receptor played an important role in the regulation of elastin distribution and artery maturation in large vessels^57, 58^. More importantly, JAG1/NOTCH3 ligand-receptor pairs were crucial for coronary arterial maturation during heart development, and the activation was induced by the appearance of blood flow^59^. Here, we discovered the role of JAG1*-*NOTCH3 signaling in regulating elastogenesis from various aspects, which reinforced the central role of VECs in regulating valvular remodeling through flow-induced Notch activation. Additionally, JAG1-Notch signaling between endothelial and adjacent smooth muscle cells in mouse ductus arteriosus and descending aorta was reported to be critical in maintaining the contractile phenotype of smooth muscle cells^60^, which is consistent with our findings showing that JAG1 secreted from UDVECs promoted the contractility of Elastin-VICs.

The importance of APOE-mediated Notch signaling was further strengthened by the loss of APOE and Notch in PVs with elastin fragmentation from individuals with PS. PV of PS individual exhibited classic myxomatous changes with pronounced elastin fragmentation^28, 31^. Interestingly, we confirmed that *JAG1* expression was reduced in UDVECs of PS valves, and as a result, *JAG1-NOTCH3* interaction between UDVEC-VIC was decreased. Although a causal relationship between APOE deficiency and valvular defects was not determined in this study, it is possible that the loss of APOE in Elastin-VICs, JAG1 inhibition in UDVECs, coupled with disturbance to blood flow, synergistically contribute to the overall decrease of Notch activation in PS Elastin-VICs, hence impairing elastogenesis during valve remodeling.

We also acknowledged several limitations in our study. The transcriptomic profile of week 15 human fetal valves used this study only represents a single time point. Although we validated the presence of Elastin-VICs in W15-17 human fetal valves via immunofluorescence and RNA *in situ* hybridization and by cross-referencing to Cui et al.’s W22-25 human fetal valve dataset, these time points only constitute a small window of the entire valve remodeling process. Another limitation is the primary human fetal VICs utilized for the *in vitro* studies. It is unclear as to whether these cells were able to fully recapitulate the transcriptomic signatures of Elastin-VICs once removed from the tissue microenvironment. Optimization of VIC culture conditions may be required to enrich the Elastin-producing VIC populations *in vitro* for downstream molecular and functional analyses. Alternatively, induced pluripotent stem cell-derived VICs (iPSC-VICs) may be an option to obtain Elastin-VICs^61, 62^. Additional work needs to be considered to determine if current iPSC-VIC differentiation protocols can generate various VIC subtypes. Nonetheless, for the purposes of this study, we were able to use primary human fetal VICs to validate the function of APOE and shear stress-induced Notch activation and elastogenesis. Lastly, due to the limit accessibility of heart valve samples from completely healthy donors, the pulmonary valves used as controls, although structurally normal, were derived from patients with coexisting cardiac conditions associated with different loading conditions and in some cases, pathology affecting other valves such as aortic valves and mitral valves.

## Supporting information

Supplement Methods, Supplement Tables S1-S2, Supplement Figures S1-S9, Reference

## Acknowledgment

The authors greatly appreciate Michelle Faust, Lindsay Fist and Olivia Croweak from Heart Institute Biorepository (HIBR), Cincinnati Children’s Hospital Medical Center (CCHMC) for collecting and providing human pulmonary valve samples; Betsy DiPasquale and Jaime Reuss from Pathology Core, the Discover Together Biobank, CCHMC for support to the study; Dr. Matt Kofron from Confocal Imaging Core (CIC), CCHMC, and Joseph Kitzmiller from Division of Pulmonary Biology, CCHMC for providing access and assistance to the confocal microscope and image processing. Author contributions: Z.L, Y.L and Z.Y contributed equally to this work. Z.L, Z.Y, Y.M, M.Gu conceived and designed experiments. Y.M collected and dissected human samples. Z.L, Y.L performed the bioinformatic analysis supervised by M.Guo. Y.W.C performed prenatal based immunofluorescent analysis. Z.L, Z.Y performed postnatal valve immunofluorescent analysis supervised by D.W.. Z.Y and Z.L performed and analyzed cell-based experiments. Z.L, Z.Y. N.P, Y.M, M.Gu wrote the manuscript with the contributions from all other authors. M.Gu, Y.M oversaw the project. All authors read and approved the manuscript.

## List of Supplemental Materials

Supplemental Methods Tables S1-S2

Figure S1-S9

Reference 1-8

## Reference

1. Aikawa E, Whittaker P, Farber M, Mendelson K, Padera RF, Aikawa M, et al. Human semilunar cardiac valve remodeling by activated cells from fetus to adult: Implications for postnatal adaptation, pathology, and tissue engineering. Circulation. 2006;113:1344–1352

2. Hinton RB, Jr., Lincoln J, Deutsch GH, Osinska H, Manning PB, Benson DW, et al. Extracellular matrix remodeling and organization in developing and diseased aortic valves. Circ Res. 2006;98:1431–1438

3. Votteler M, Berrio DA, Horke A, Sabatier L, Reinhardt DP, Nsair A, et al. Elastogenesis at the onset of human cardiac valve development. Development. 2013;140:2345–2353

4. Lindsey SE, Butcher JT, Yalcin HC. Mechanical regulation of cardiac development. Front Physiol. 2014;5:318

5. Stuart AG, Williams A. Marfan’s syndrome and the heart. Arch Dis Child. 2007;92:351–356

6. Waller BF, Howard J, Fess S. Pathology of pulmonic valve stenosis and pure regurgitation. Clin Cardiol. 1995;18:45–50

7. Krishnan A, Samtani R, Dhanantwari P, Lee E, Yamada S, Shiota K, et al. A detailed comparison of mouse and human cardiac development. Pediatr Res. 2014;76:500–507

8. Liu AC, Joag VR, Gotlieb AI. The emerging role of valve interstitial cell phenotypes in regulating heart valve pathobiology. Am J Pathol. 2007;171:1407–1418

9. Goddard LM, Duchemin AL, Ramalingan H, Wu B, Chen M, Bamezai S, et al. Hemodynamic forces sculpt developing heart valves through a klf2-wnt9b paracrine signaling axis. Dev Cell. 2017;43:274–289 e275

10. Hogers B, DeRuiter MC, Gittenberger-de Groot AC, Poelmann RE. Extraembryonic venous obstructions lead to cardiovascular malformations and can be embryolethal. Cardiovasc Res. 1999;41:87–99

11. Hove JR, Koster RW, Forouhar AS, Acevedo-Bolton G, Fraser SE, Gharib M. Intracardiac fluid forces are an essential epigenetic factor for embryonic cardiogenesis. Nature. 2003;421:172–177

12. Vermot J, Forouhar AS, Liebling M, Wu D, Plummer D, Gharib M, et al. Reversing blood flows act through klf2a to ensure normal valvulogenesis in the developing heart. PLoS Biol. 2009;7:e1000246

13. Yalcin HC, Shekhar A, Nishimura N, Rane AA, Schaffer CB, Butcher JT. Two-photon microscopy-guided femtosecond-laser photoablation of avian cardiogenesis: Noninvasive creation of localized heart defects. Am J Physiol Heart Circ Physiol. 2010;299:H1728–1735

14. Cui Y, Zheng Y, Liu X, Yan L, Fan X, Yong J, et al. Single-cell transcriptome analysis maps the developmental track of the human heart. Cell Rep. 2019;26:1934–1950 e1935

15. Leshem RS, Baker SM, Mallen J, Wang L, Dark J, Bamforth S, et al. A cell atlas of the human outflow tract of the heart and its adult derivatives. bioRxiv. 2023:2023.2004.2005.535627

16. Hulin A, Hortells L, Gomez-Stallons MV, O’Donnell A, Chetal K, Adam M, et al. Maturation of heart valve cell populations during postnatal remodeling. Development. 2019;146

17. Stuart T, Butler A, Hoffman P, Hafemeister C, Papalexi E, Mauck WM, 3rd, et al. Comprehensive integration of single-cell data. Cell. 2019;177:1888–1902 e1821

18. Hinton RB, Yutzey KE. Heart valve structure and function in development and disease. Annu Rev Physiol. 2011;73:29–46

19. Gow A, Southwood CM, Li JS, Pariali M, Riordan GP, Brodie SE, et al. Cns myelin and sertoli cell tight junction strands are absent in osp/claudin-11 null mice. Cell. 1999;99:649–659

20. Shimomura Y, Agalliu D, Vonica A, Luria V, Wajid M, Baumer A, et al. Apcdd1 is a novel wnt inhibitor mutated in hereditary hypotrichosis simplex. Nature. 2010;464:1043–1047

21. Huang Y, Mahley RW. Apolipoprotein e: Structure and function in lipid metabolism, neurobiology, and alzheimer’s diseases. Neurobiol Dis. 2014;72 Pt A:3-12

22. Murakami M, Sato H, Miki Y, Yamamoto K, Taketomi Y. A new era of secreted phospholipase a(2). J Lipid Res. 2015;56:1248–1261

23. Tanaka K, Sata M, Fukuda D, Suematsu Y, Motomura N, Takamoto S, et al. Age-associated aortic stenosis in apolipoprotein e-deficient mice. J Am Coll Cardiol. 2005;46:134–141

24. Aikawa E, Nahrendorf M, Sosnovik D, Lok VM, Jaffer FA, Aikawa M, et al. Multimodality molecular imaging identifies proteolytic and osteogenic activities in early aortic valve disease. Circulation. 2007;115:377–386

25. Hartley CJ, Reddy AK, Madala S, Martin-McNulty B, Vergona R, Sullivan ME, et al. Hemodynamic changes in apolipoprotein e-knockout mice. Am J Physiol Heart Circ Physiol. 2000;279:H2326–2334

26. Milewicz DM, Trybus KM, Guo DC, Sweeney HL, Regalado E, Kamm K, et al. Altered smooth muscle cell force generation as a driver of thoracic aortic aneurysms and dissections. Arterioscler Thromb Vasc Biol. 2017;37:26–34

27. Karimi A, Milewicz DM. Structure of the elastin-contractile units in the thoracic aorta and how genes that cause thoracic aortic aneurysms and dissections disrupt this structure. Can J Cardiol. 2016;32:26–34

28. Grande-Allen KJ, Griffin BP, Ratliff NB, Cosgrove DM, Vesely I. Glycosaminoglycan profiles of myxomatous mitral leaflets and chordae parallel the severity of mechanical alterations. J Am Coll Cardiol. 2003;42:271–277

29. Kim AJ, Xu N, Umeyama K, Hulin A, Ponny SR, Vagnozzi RJ, et al. Deficiency of circulating monocytes ameliorates the progression of myxomatous valve degeneration in marfan syndrome. Circulation. 2020;141:132–146

30. Saef JM, Ghobrial J. Valvular heart disease in congenital heart disease: A narrative review. Cardiovasc Diagn Ther. 2021;11:818–839

31. Escalon JG, Browne LP, Bang TJ, Restrepo CS, Ocazionez D, Vargas D. Congenital anomalies of the pulmonary arteries: An imaging overview. Br J Radiol. 2019;92:20180185

32. Allen WM, Matloff JM, Fishbein MC. Myxoid degeneration of the aortic valve and isolated severe aortic regurgitation. Am J Cardiol. 1985;55:439–444

33. Jin S, Guerrero-Juarez CF, Zhang L, Chang I, Ramos R, Kuan CH, et al. Inference and analysis of cell-cell communication using cellchat. Nat Commun. 2021;12:1088

34. Bowers SL, Banerjee I, Baudino TA. The extracellular matrix: At the center of it all. J Mol Cell Cardiol. 2010;48:474–482

35. Henneman P, van der Sman-de Beer F, Moghaddam PH, Huijts P, Stalenhoef AF, Kastelein JJ, et al. The expression of type iii hyperlipoproteinemia: Involvement of lipolysis genes. Eur J Hum Genet. 2009;17:620–628

36. Sakai LY, Keene DR, Renard M, De Backer J. Fbn1: The disease-causing gene for marfan syndrome and other genetic disorders. Gene. 2016;591:279–291

37. Rifkin DB, Rifkin WJ, Zilberberg L. Ltbps in biology and medicine: Ltbp diseases. Matrix Biol. 2018;71–72:90-99

38. Maki JM, Rasanen J, Tikkanen H, Sormunen R, Makikallio K, Kivirikko KI, et al. Inactivation of the lysyl oxidase gene lox leads to aortic aneurysms, cardiovascular dysfunction, and perinatal death in mice. Circulation. 2002;106:2503–2509

39. Watanabe K, Taketomi Y, Miki Y, Kugiyama K, Murakami M. Group v secreted phospholipase a(2) plays a protective role against aortic dissection. J Biol Chem. 2020;295:10092–10111

40. Obniski R, Sieber M, Spradling AC. Dietary lipids modulate notch signaling and influence adult intestinal development and metabolism in drosophila. Dev Cell. 2018;47:98–111 e115

41. Theendakara V, Peters-Libeu CA, Spilman P, Poksay KS, Bredesen DE, Rao RV. Direct transcriptional effects of apolipoprotein e. J Neurosci. 2016;36:685–700

42. Foraker J, Millard SP, Leong L, Thomson Z, Chen S, Keene CD, et al. The apoe gene is differentially methylated in alzheimer’s disease. J Alzheimers Dis. 2015;48:745–755

43. Sen A, Nelson TJ, Alkon DL. Apoe4 and abeta oligomers reduce bdnf expression via hdac nuclear translocation. J Neurosci. 2015;35:7538–7551

44. Ferrante F, Giaimo BD, Bartkuhn M, Zimmermann T, Close V, Mertens D, et al. Hdac3 functions as a positive regulator in notch signal transduction. Nucleic Acids Res. 2020;48:3496–3512

45. Wunnemann F, Ta-Shma A, Preuss C, Leclerc S, van Vliet PP, Oneglia A, et al. Loss of adamts19 causes progressive non-syndromic heart valve disease. Nat Genet. 2020;52:40–47

46. Oikonomopoulos A, Sereti KI, Conyers F, Bauer M, Liao A, Guan J, et al. Wnt signaling exerts an antiproliferative effect on adult cardiac progenitor cells through igfbp3. Circ Res. 2011;109:1363–1374

47. Hulin A, Moore V, James JM, Yutzey KE. Loss of axin2 results in impaired heart valve maturation and subsequent myxomatous valve disease. Cardiovasc Res. 2017;113:40–51

48. Miao Y, Tian L, Martin M, Paige SL, Galdos FX, Li J, et al. Intrinsic endocardial defects contribute to hypoplastic left heart syndrome. Cell Stem Cell. 2020;27:574–589 e578

49. Yu Z, Zhou X, Liu Z, Pastrana-Gomez V, Liu Y, Guo M, et al. Kmt2d-notch mediates coronary abnormalities in hypoplastic left heart syndrome. Circ Res. 2022;131:280–282

50. Yu Z, Liu Z, Ravichandran V, Lami B, Gu M. Endocardium in hypoplastic left heart syndrome: Implications from in vitro study. J Cardiovasc Dev Dis. 2022;9

51. Qu X, Violette K, Sewell-Loftin MK, Soslow J, Saint-Jean L, Hinton RB, et al. Loss of flow responsive tie1 results in impaired aortic valve remodeling. Dev Biol. 2019;455:73–84

52. Theodoris CV, Li M, White MP, Liu L, He D, Pollard KS, et al. Human disease modeling reveals integrated transcriptional and epigenetic mechanisms of notch1 haploinsufficiency. Cell. 2015;160:1072–1086

53. Mack JJ, Mosqueiro TS, Archer BJ, Jones WM, Sunshine H, Faas GC, et al. Notch1 is a mechanosensor in adult arteries. Nat Commun. 2017;8:1620

54. Masumura T, Yamamoto K, Shimizu N, Obi S, Ando J. Shear stress increases expression of the arterial endothelial marker ephrinb2 in murine es cells via the vegf-notch signaling pathways. Arterioscler Thromb Vasc Biol. 2009;29:2125–2131

55. Turnpenny PD, Ellard S. Alagille syndrome: Pathogenesis, diagnosis and management. Eur J Hum Genet. 2012;20:251–257

56. Hofmann JJ, Briot A, Enciso J, Zovein AC, Ren S, Zhang ZW, et al. Endothelial deletion of murine jag1 leads to valve calcification and congenital heart defects associated with alagille syndrome. Development. 2012;139:4449–4460

57. Krebs LT, Xue Y, Norton CR, Sundberg JP, Beatus P, Lendahl U, et al. Characterization of notch3-deficient mice: Normal embryonic development and absence of genetic interactions with a notch1 mutation. Genesis. 2003;37:139–143

58. Domenga V, Fardoux P, Lacombe P, Monet M, Maciazek J, Krebs LT, et al. Notch3 is required for arterial identity and maturation of vascular smooth muscle cells. Genes Dev. 2004;18:2730–2735

59. Volz KS, Jacobs AH, Chen HI, Poduri A, McKay AS, Riordan DP, et al. Pericytes are progenitors for coronary artery smooth muscle. Elife. 2015;4

60. Feng X, Krebs LT, Gridley T. Patent ductus arteriosus in mice with smooth muscle-specific jag1 deletion. Development. 2010;137:4191–4199

61. Neri T, Hiriart E, van Vliet PP, Faure E, Norris RA, Farhat B, et al. Human pre-valvular endocardial cells derived from pluripotent stem cells recapitulate cardiac pathophysiological valvulogenesis. Nat Commun. 2019;10:1929

62. Cheng L, Xie M, Qiao W, Song Y, Zhang Y, Geng Y, et al. Generation and characterization of cardiac valve endothelial-like cells from human pluripotent stem cells. Commun Biol. 2021;4:1039

