## Supplement Methods, Supplement Tables S1-S2, Supplement Figures S1-S9, Reference for "APOE-NOTCH Axis Governs Elastogenesis During Human Cardiac Valve Remodeling"

**Supplemental Material**

#### Supplemental Methods

##### Human Heart Tissue Section

Postnatal pulmonary valve tissue specimens were acquired from patients diagnosed with Tetralogy of Fallot (TOF) or other types of non-pulmonary valve related heart disease (control). Both fetal and postnatal heart tissues were fixed in 4% paraformaldehyde overnight at 4°C, dehydrated through a graded ethanol series, cleared in xylene, embedded in paraffin, and sectioned at 5μm thickness.

##### Histology and Immunostaining

Movat's Pentachrome Staining

Paraffin tissue sections were dewaxed in 2 changes of xylene and dehydrated 3 changes of absolute alcohol. Verhoeff's elastic stain reagent was stained for 15min for elastin staining, 1% alcian blue solution was stained for 15min for acidic mucopolysaccharides staining, Crocein scarlet-acid fuchsin was stained for 2min for muscle staining, Collagen and reticulin fibers are unstained by 2 changes of 5% phosphotungstic acid for 2min/change and stained in yellow by alcoholic saffron solution for 15min. The slides were then dehydrated by 3 changes of absolute alcohol and 3 changes of xylene, and were mounted by Cytoseal mounting media (Thermo Scientific, Ref#8312-4) The procedure of Movat's Pentachrome staining was performed strictly following manufacturer's protocols (American MasterTech).

Immunofluorescence Staining

Cells were fixed in 4% paraformaldehyde for 15 min and blocked with 5% donkey serum (Jackson lab, Cat#017-000-121) diluted in PBST (0.1% Triton-X) for 1 hour. Cells were stained with primary antibodies diluted in 5% donkey serum at 4°C overnight. The human healthy control heart and stenotic valve tissue paraffin sections were dewaxed, rehydrated and performed antigen retrieval using Tris-based buffer (Vector Laboratories). The sections were then blocked with 5% Donkey serum in PBST for 1 hour. Primary antibodies: CD31 (1:100, R&D, AF806), Elastin (1:100, Millipore, MAB2503), APOE (1:100, Abcam, ab183597), PROX1 (1:100, R&D, AF2727), EMILIN1 (1:100, Sigma, HPA002822), LOXL1 (1:100, Novus Biologicals, H00004016-D01P), α-SMA (1:100, Sigma, A2547), CD55 (1:100, R&D, AF2009), FOXC2 (1:100, Proteintech, 23066-1-AP) and NOTCH1 (1:100, Abcam, ab52627). AlexaFluor-conjugated secondary antibody (Life Technologies) was then used and co-stained with DAPI (Vector Laboratories). Imaging was captured with a confocal microscope (Nikon) and analyzed with NIS-Element Analysis software.

RNA *in-situ* Hybridization

For *in-situ* mRNA detection, a proprietary high sensitivity RNA amplification and detection technology (RNA-SCOPE, Advanced Cell Diagnostics) was performed. Paraffin-embedded heart sections (5μm) were processed for *in-situ* detection using proprietary probes: *CD31* (443471), *APOE* (433091), *PTGDS* (431471), *JAG1* (546181) and *NOTCH3* (558991), and *PROX1* (530241), the multiplex fluorescent assay v2 detection kit (Cat# 323110) was used according to manufacturer's instructions. Briefly, Paraffin section slides were baked in a dry oven for 1 hour at 60°C. Sections were deparaffinized in 2 changes of xylene for 5min/changes and 1 change of 100% EtOH for 2min, then the sections were air-dried completely for around 10min. The RNAscope® Hydrogen Peroxide were applied on the sections, the slides were incubated in HybEZ™ II Oven at 40°C for 10min. After washing in distill water, the slides were put into the 1L beaker with boiled co-detection target retrieval solution for 15min. After cooling down, sections were covered by drops of RNAscope Protease Plus and incubated in HybEZ™ II Oven at 40°C for 30min. After the Protease Plus treatment and water wash, drops of probe mixture (volume ratio C1:C2:C3=50:1:1) were applied on the sections, the sections were incubated for 2hour at 40°C. The slides were then washed and transferred into 5X SSC overnight at RT. On the second day, sections were covered by AMP and incubated at 40°C for 10min. This step was repeated one by one for each AMP (AMP1/2/3) if multiple AMPs were applied. Then, drops of HRP-C1 were applied at 40°C for 15min and washed, which was followed by staining of diluted Opal 520 (diluted in TSA buffer 1:250). This step was repeated if AMP2 (HRP-C2, Opal 570 diluted in 1:500) and AMP3 (HRP-C3, Opal 690 diluted in 1:500) were used. Last, the sections were counterstained DAPI and mounted by ProLong Gold Antifade Mountant (Invitrogen, P36934) and the slides were dried overnight in the dark at 4°C.

**Isolation of Human Fetal Heart Valve Endothelial Cells and Interstitial Cells**

Human fetal hearts were obtained from aborted fetuses (Males and females, gestational week 13-19) with parental consent at the University of Washington BDRL and preserved in Belzer UW® Cold Storage Solution (Thermo Fisher) for overnight shipping. The heart was kept on ice throughout processing and 1X HBSS (GIBCO) was used to remove excess blood cells. Aortic, pulmonary, mitral, and tricuspid valves were collected. Digestion buffer was prepared in DMEM medium with each component as the following final concentration: Liberase TM, 0.5 mg/ml; DNase I, 20μg/ml; HEPEs, 10mM.

For valve endothelial cell (VEC) isolation, the valve tissue was immersed in approximately 500μL of digestion buffer. After digesting for 5 min at 37°C, the tissue was washed with EGM-2 medium (Lonza) prior to centrifugation to pellet the cells. Then the cells were separated through a 40μm cell strainer and sorted using a MACS sorter with a CD144 antibody (Miltenyi Biotec). EC populations were tested for CD144+ by flow cytometry and purity by quantitative PCR. Sorted cells were resuspended with EGM-2 containing an additional 50ng/ml VEGF and 10ng/ml bFGF and seeded on a collagen I (Sigma)-coated plate. The remaining CD144- valve interstitial cells (VICs) were resuspended with FGM-3 Cardiac Fibroblast Growth Medium (Lonza) and cultured in culture dish. The human fetal VECs (Passage 3-7) and VICs (Passage 3-10) were utilized for downstream experiments. Cell identities were validated through qPCR detecting of marker genes (**Figure S6M**), immunofluorescent staining of cell-type specific markers (**Figure S6N**), and flow cytometry analysis (**Figure S6O**).

**Cell Culture**

Human VICs were kept in FGM3 medium (Lonza, CC-4526) in culture dish or plate. Jag1 ligand treatment was established as described before[1]. Briefly, culture plates were coated with Jag1 (15μg/ml) (R&D system, Cat#1277-JG) in PBS at 4°C overnight. VICs were seeded in Jag1 coated plate for 48 hours before analysis. Human VECs were cultured in plate in EGM2 medium (Lonza, CC-3162), which was coated with 0.2% gelatin solution (Sigma, G1393) for 30 min in 37°C. Cells were routinely maintained in 95% air and 5% $CO_2$ at 37°C.

**Co-culture of VICs and VECs**

All co-cultures were performed using 0.4μm pore 6-well plate cell culture inserts (Cellqart) coated with 0.2% gelatin solution for 30min in 37°C. Monoculture was established by seeding EC on the bottom of the insert, and contact co-culture was established as previously described[2]. Briefly, VICs suspended in FGM3 medium were seeded on the opposite side of an insert. Two hours after seeding, the inserts were flipped, and EC were seeded on the bottom of the inserts and cultivated in EC medium. 1ml EGM2 was added in the upper inserts for EC culturing, 2ml FGM3 was added in the lower well of a 6-well plate for VIC culturing.

**Quantification of Tropo-elastin and Mature Elastin**

The Fastin Elastin Assay (Biocolor Life Science Assays, F2000) was used to measure the insoluble (mature) elastin content and intracellular soluble tropo-elastin. VICs treated in different conditions were cultured in T25 flask till 90% confluency. Cells were digested with 0.25 M oxalic acid at 100 °C for 1 hour for twice. An equal volume of elastin precipitating reagent was added to the total two extractions of digested solution for 15min. Then, elastin was precipitated by centrifuge at 13000 rpm for 10min. The supernatant was then used for protein quantification by bicinchoninic acid assay. The assay was then performed according to the manufacturer's instructions, and the elastin content was calculated based on the standard curve and normalized to the total protein content.

**Orbital Shear Stress**

EC monoculture or EC-VIC coculture were established on the cellqart insert in a 6-well plate. An orbital unidirectional flow was applied to confluent ECs by placing the plate on an orbital shaker (Talboys) at 250rpm. The laminar shear stress exerted on EC was calculated as 8.52 dynes/cm$^2$ based on the previous described formula $\tau_w = a\sqrt{\rho\eta\omega^3}$[3]. The orbital flow system was kept in a 5% $CO_2$ humidified incubator at 37°C. Morphological changes were observed under a microscope at an original magnification of x10. 72 hours post flow induction, VECs and VICs were digested separately from two sides of the insert membrane for RNA extraction. The insert membrane was cut off for following immunostaining.

**Quantitative Reverse-transcription PCR**

Our detailed protocol was previously published[4]. Briefly, total RNA was extracted, purified, and quantified for reverse transcription using High-Capacity RNA to cDNA Kit (Applied Biosystems) according to the manufacturer's instructions. qPCR was carried out using 5ng cDNA and 6µL SYBR green master mix (Applied Biosystems). Primers were listed in Supplemental Table 2. Each measurement was performed in triplicate.

**Flow Cytometry Analysis**

Human primary interstitial cells (VICs) and valve endothelial cells (VECs) were dissociated and resuspended into single cells in cell staining buffer (BioLegend, Cat#420201) after being filtered through 40um cell strainer. Single cells were aliquoted in cell staining buffer (between 100,000 and 125,000 cells were used per cell sample) and treated with unstained controls or antigen-specific antibodies (CD31, R&D, AF806 R&D) for 20 min at 4ºC. Cells were washed and stained with secondary fluorescent antibody in cell staining buffer for 20min in the dark. (Donkey anti-sheep-488, 1:200, Jackson lab, cat#713-545-147). After twice washes, cells were fixed in fixation buffer (BioLegend, Cat. No. 420801) for 20min in the dark and permeabilized by resuspending fixed cells in intracellular staining perm wash buffer (BioLegend, Cat#421002) and centrifuged at 350xg for 5min twice. Intracellular staining was performed by resuspending cells in intracellular staining perm wash buffer and adding fluorophore-conjugated antibodies at 1:100 diluted ratio (COL1a1,

Southern Biotech 1310-01). Intracellular staining was performed for 20min in the dark. Last, cells were washed twice and resuspended in cell staining buffer and filtered into flow test tube (Falcon, 352235) for flow cytometry analysis. Flow cytometry was performed using analytical cytometer (BD/LSR Fortessa). Data analysis was performed with FlowJo_V10. Ten thousand events were collected in each run.

**siRNA Transfection**

siRNA transfection was carried out according to manufacturer's instruction. ON-TARGET plus scramble, APOE siRNA (Horizon Dharmacon) were transfected at 25nM into sub-confluent early passage VICs using Lipofectamine RNAiMax (Invitrogen). 48h after transfection, cells were either fixed for staining, subjected to RNA extraction or downstream contractility assay.

**Gel Contraction Assay**

Collagen gel contraction assays were performed according to the manufacturer's instructions (Cell Biolabs, CBA-201). Briefly, cells were pretreated with APOE siRNA or 10μm NOTCH inhibitor (L-685,458, TOCRIS Cat#2627) or PBS for 24h. The cells were then collected from plates and suspended at a density of $4.0 \times 10^5$ cells per 100μL of medium with or without 15ug/ml Jag1 ligands to which 400μL of neutralized collagen solution were added. The resultant mixture was added to one well of a 24-well culture plate and allowed to gel for 2 hours at 37 °C. After polymerization, 1mL of FGM3 culture medium was added on top of the gel lattice and incubated for 48 hours. The stress was then released by running a sterile pipette tip along the sides of the well. 24 hours after the stress was released, the culture dish was scanned, and the area of the collagen gel was measured using ImageJ software version 2.3.0/1.53f (NIH, Bethesda, MD; http://imagej.nih.gov/ij)

**scRNA-seq Data Pre-processing**

For pre-processing of the single-cell RNA-seq data, we utilized Chromium Single Cell Software Suite v1.1 from 10X Genomics. This software suite facilitated sample de-multiplexing, barcode processing, and single-cell 3' gene counting. Sample de-multiplexing was performed based on the 8-bp sample index read from the Illumina BCL output folder. Subsequently, FASTQ files were generated for the single end reads and sample index using Illumina bcl2fastq. The reads were then mapped to the human reference genome (GRCh38) using STAR. Confidently mapped reads were delivered in a BAM file. Gene-barcode matrices were generated using the Chromium cellular barcodes, and filtered gene-barcode matrices containing only cellular barcodes in MEX format were used for downstream analysis.

**Data Analysis of scRNA-seq**

Only genes expressed in at least three cells and cells with a detected gene count between 1,000 and 12,000 were retained. Additionally, cells with a high percentage of mitochondrial genes (>10%) and a high scrublet score (>0.3) were filtered out. The Seurat V3 data integration pipeline was used to batch correct the data through the canonical correlation analysis (CCA) method. According to a benchmark comparison study conducted by Tran and colleagues[5], Seurat 3 CCA was identified as one of the top three preferred batch integration techniques for this type of data. The R package SCTransform[6] was used to normalize gene expression for each cell by fitting the Gamma-Poisson generalized linear model. The resulting log-transformed, normalized single-cell expression values were used for visualizations and differential expression tests. Statistically significant principal components were determined by a resampling test and were retained for the Uniform Manifold Approximation and Projection (UMAP) analysis. Differential expression analysis among clusters was conducted using a likelihood-ratio test, comparing cells within each cluster against all other cells. Gene A was defined as a biomarker for cluster X if it was detected in at least 25% of cells, had an adjusted p-value less than 0.05, and had a log e fold change of at least 0.25 between cells of cluster X and all other cells. These analyses were performed using the Seurat package v4.0. DEGs were analyzed for GO terms and KEGG pathways enrichment by using KOBAS[7]. A significance threshold of FDR < 0.05 was used during the enrichment analysis to identify significant results.

**Projection of scRNA-seq Data to Reference Data**

The Seurat R package was used to project single-cell transcriptome data from mouse and human valves onto our human valve data. Anchors between the two datasets were calculated using the top 50 principal components from the reference data through the FindTransferAnchors function. The TransferData function was used to predict the cell type of each cell in the query data based on the anchors within 50 dimensions for the anchor weighting procedure. The query data were then projected to the reference data using MapQuery (reference.reduction as 'pca' and reduction.model as 'umap'), resulting in a UMAP plot of combined both mouse and human data.

**Cell-cell Communication Analysis**

To analyze cell-cell communication between VIC and VEC, we utilized the CellChat R package[8]. The normalized counts were loaded into CellChat and standard preprocessing steps were applied, including the functions identifyOverExpressedGenes, identifyOverExpressedInteractions, and projectData with standard parameters. A pre-compiled network of human ligand-receptor interactions was used as a priori information. We calculated the potential ligand-receptor interactions between VIC and VEC using the functions computeCommunProb, computeCommunProbPathway, and aggregateNet with standard parameters. Herein, we set Ligand-Receptor pairs with probability higher than 0.001 and P value smaller than 0.05 as significant.

#### Table S1. Demographic information for human valve samples

| Sample name | Category | Specimen | Age | Gender | Race | Diagnosis, Pathology |
|---|---|---|---|---|---|---|
| TOF1 | "Disease" | Pulmonary valve | 10 months | Female | White | 2 short thick leaflets of cardiac valvular tissue with prominent increase in myxoid ground substance. Prominent myxoid change suggesting valvular dysplasia. |
| TOF2 | "Disease" | Pulmonary valve | 0-1 year | Male | White | Pieces of dysplastic pulmonary valve. Clinical history of Tetralogy of Fallot. |
| TOF3 | "Disease" | Pulmonary valve | 5 months | Male | White | Tetralogy of Fallot, Pulmonary Stenosis with dysplastic valve structure. |
| Control-1 | "Control" | Pulmonary valve | 9 months | Male | White | HLHS |
| Control-2 | "Control" | Pulmonary valve | 7 years | Male | White | HLHS |
| Control-3 | "Control" | Pulmonary valve | 10-year 7 month | Male | White | HCM |

**Table S2. qPCR primer sequence**

| Primer names | Sequence |
|---|---|
| Hu-APOE-S | GGACGTCCTTCCCCAGGA |
| Hu-APOE-AS | GAATGTGACCAGCAACGCAG |
| Hu-ELN-S | TTCCCCGCAGTTACCTTTCC |
| Hu-ELN-AS | CTAAGCCACCAACTCCTGGG |
| Hu-FBN1-S | GCGGAAATCAGTGTATTGTCCC |
| Hu-FBN1-AS | CAGTGTTGTATGGATCTGGAGC |
| Hu-FBN2-S | TGAGTGCAGCATCATTCCTGG |
| Hu-FBN2-AS | AGAAACACATGCCTGTTCTCTG |
| Hu-EMILIN1-S | CAGCCTCTACACAGGTTCCAG |
| Hu-EMILIN1-AS | CACGTAGGCACACCAGTTCC |
| Hu-LOX-S | GCCGACCAAGATATTCCTGGG |
| Hu-LOX-AS | GCAGGTCATAGTGGCTAAACTC |
| Hu-LOXL1-S | CCACTACGACCTACTGGATGC |
| Hu-LOXL1-AS | GTTGCCGAAGTCACAGGTG |
| Hu-FBLN2-S | CAGGTGGCCTCTAACACCATC |
| Hu-FBLN2-AS | CTGCTTGCAGGGTCCATTGT |
| Hu-FBLN1-S | AGAGCTGCGAGTACAGCCT |
| Hu-FBLN1-AS | CGACATCCAAATCTCCGGTCT |
| Hu-LTBP4-S | CTGCCCATTCTGCGGAACAT |
| Hu-LTBP4-AS | GCCGAGTAGTGGTAACCAGG |
| Hu-LTBP2-S | ACGCGGAGTGTGTGAATACC |
| Hu-LTBP2-AS | GCAGCATGGAGATTGCCTTG |
| Hu-LTBP1-S | CTGACGGCCACGAACTTCC |
| Hu-LTBP1-AS | GCACTGACATTTGTCCCTTGA |
| Hu-TIMP2-S | GCTGCGAGTGCAAGATCAC |
| Hu-TIMP2-AS | TGGTGCCCGTTGATGTTCTTC |
| Hu-MMP14-S | GGCTACAGCAATATGGCTACC |
| Hu-MMP14-AS | GATGGCCGCTGAGAGTGAC |
| Hu-TIMP3-S | CATGTGCAGTACATCCATACGG |
| Hu-TIMP3-AS | CATCATAGACGCGACCTGTCA |
| Hu-MYH11-S | AGATGGTTCTGAGGAGGAAACG |
| Hu-MYH11-AS | AAAACTGTAGAAAGTTGCTTATTCACT |
| Hu-ACTA2-S | GTGTTGCCCCTGAAGAGCAT |
| Hu-ACTA2-AS | GCTGGGACATTGAAAGTCTCA |
| Hu-TAGLN-S | CCGTGGAGATCCCAACTGG |
| Hu-TAGLN-AS | CCATCTGAAGGCCAATGACAT |
| Hu-SMTN-S | CCCTGGCATCCAAGCGTTT |
| Hu-SMTN-AS | CTCCACATCGTTCATGGACTC |
| Hu-KI67-S | ACGCCTGGTTACTATCAAAAGG |
| Hu-KI67-AS | CAGACCCATTTACTTGTGTTGGA |
| Hu-CDK4-S | ATGGCTACCTCTCGATATGAGC |

**Table S2. qPCR primer sequence (Continued)**

| Primer names | Sequence |
|---|---|
| Hu-CDK4-AS | CATTGGGGACTCTCACACTCT |
| Hu-CCND1-S | GCTGCGAAGTGGAAACCATC |
| Hu-CCND1-AS | CCTCCTTCTGCACACATTTGAA |
| Hu-NOTCH1-S | GAGGCGTGGCAGACTATGC |
| Hu-NOTCH1-AS | CTTGTACTCCGTCAGCGTGA |
| Hu-JAG1-S | GGGGCAACACCTTCAACCTC |
| Hu-JAG1-AS | CCAGGCGAAACTGAAAGGC |
| Hu-HES1-S | ACGTGCGAGGGCGTTAATAC |
| Hu-HES1-AS | GGGGTAGGTCATGGCATTGA |
| Hu-HEY1-S | GTTCGGCTCTAGGTTCCATGT |
| Hu-HEY1-AS | CGTCGGCGCTTCTCAATTATTC |
| Hu-HEY2-S | GCCCGCCCTTGTCAGTATC |
| Hu-HEY2-AS | CCAGGGTCGGTAAGGTTTATTG |
| Hu-HEYL-S | GGAAGAAACGCAGAGGGATCA |
| Hu-HEYL-AS | CAAGCGTCGCAATTCAGAAAG |
| Hu-DLL4-S | GTCTCCACGCCGGTATTGG |
| Hu-DLL4-AS | CAGGTGAAATTGAAGGGCAGT |
| Hu-NOS3-S | TGATGGCGAAGCGAGTGAAG |
| Hu-NOS3-AS | ACTCATCCATACACAGGACCC |
| Hu-NOTCH3-S | TGGCGACCTCACTTACGACT |
| Hu-NOTCH3-AS | CACTGGCAGTTATAGGTGTTGAC |
| Hu-DLL1-S | GACGAACACTACTACGGAGAGG |
| Hu-DLL1-AS | AGCCAGGGTTGCACACTTT |
| Hu-KLF2-S | CTACACCAAGAGTTCGCATCTG |
| Hu-KLF2-AS | CCGTGTGCTTTCGGTAGTG |
| Hu-KLF4-S | CAGCTTCACCTATCCGATCCG |
| Hu-KLF4-AS | GACTCCCTGCCATAGAGGAGG |

#### Figure S1

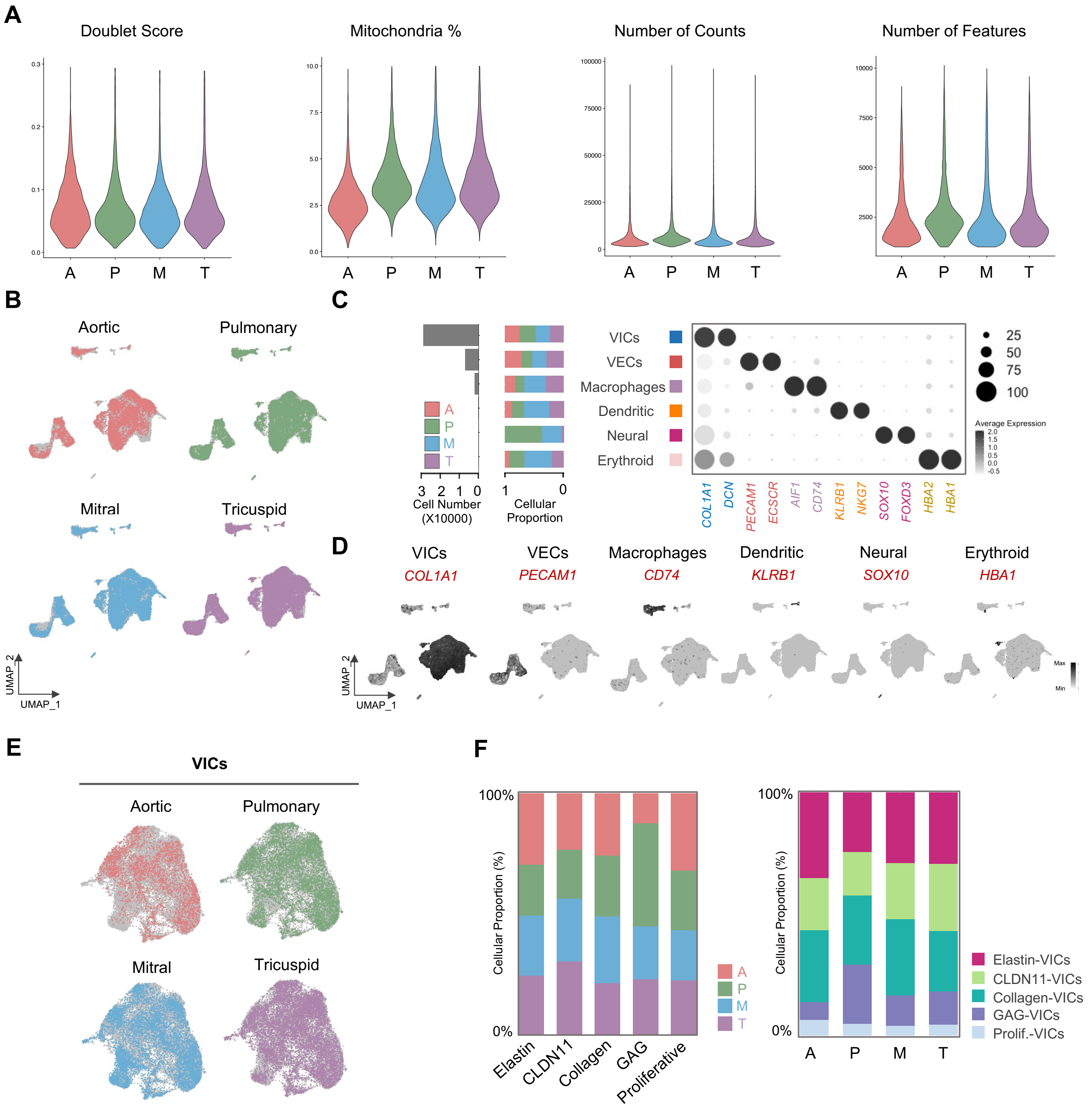


**Figure S1. scRNA-seq of Normal Human Fetal Heart Valves**

**A.** Violin plot of doublet score, mitochondria percentage, number of counts and number of features across all valves after quality control: nCount RNA < 1e5, nFeature RNA >= 1000, pMT < 10, Doublet score <0.3. A/P/M/T represent Aortic, Pulmonary, Mitral, Tricuspid valves respectively; **B.** UMAP visualization of cells from four different human fetal valves from one normal heart after integration by Canonical Correlation Analysis (CCA); **C.** Left: Cell number and proportion of each valve in each valve cell type. Right: Dot plot of representative marker genes within each valve cell type; **D.** Feature plots of representative marker genes in each valve cell type; **E.** UMAP visualization of four different human fetal valves within VICs; **F.** Left: Bar graph showing proportions of each valve within each VIC subtype. Right: Bar graph showing proportions of each VIC subtype within each valve.

#### Figure S2

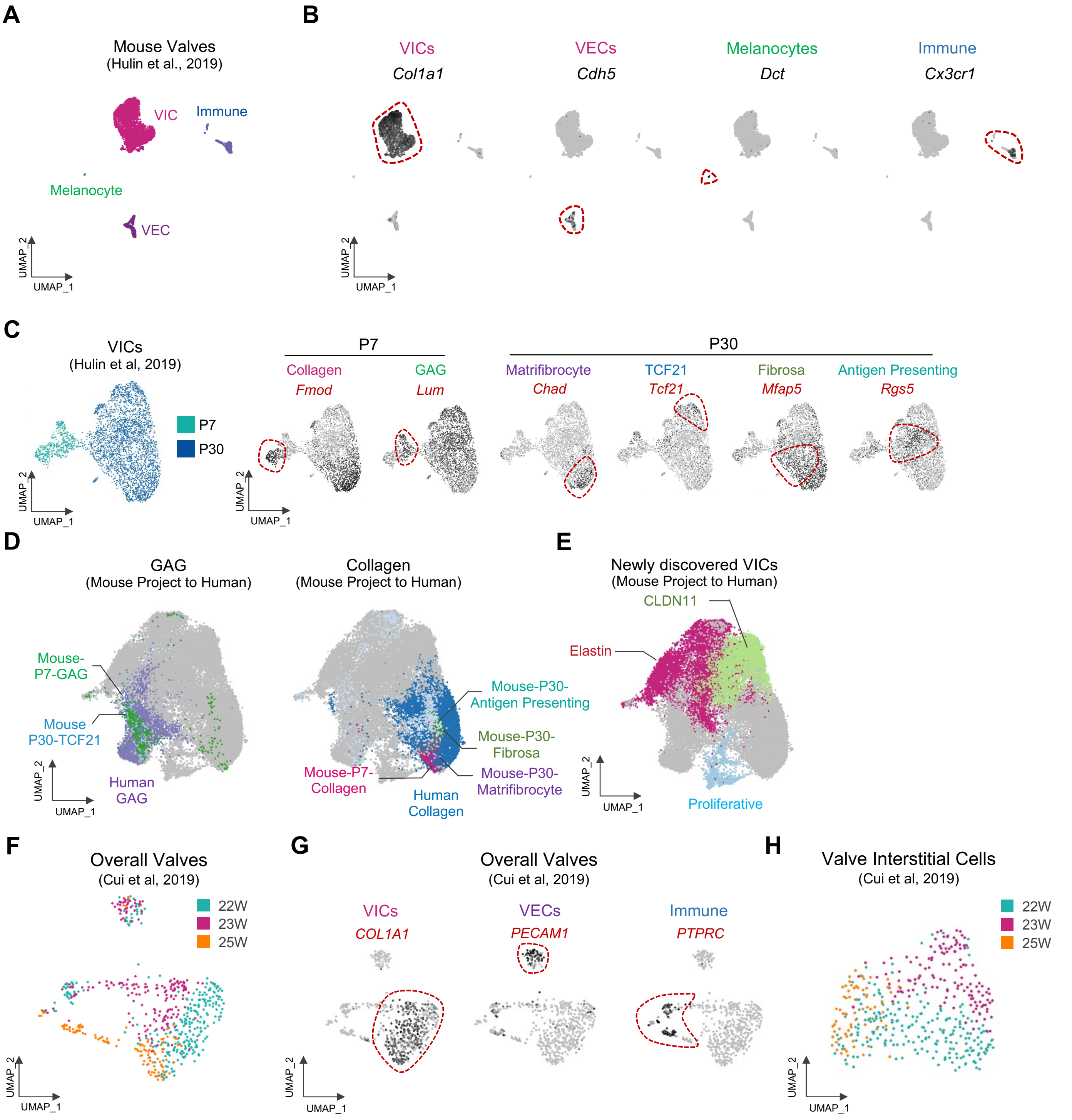


**Figure S2. Re-analysis of Public Human Fetal and Mouse Postnatal Heart Valve scRNA-seq datasets.** **A.** UMAP visualization of postnatal mouse valve cell types (Hulin et al., 2019); **B.** Feature plots of representative marker genes within each cell type within mouse valves; **C.** Development stage visualization (P7 and P30, Left) and feature plots (Right) of representative marker genes of each VIC subtype; **D.** UMAP visualization showing mouse VIC subtypes similar to human GAG-VICs (Left) and Collagen-VICs (Right) through reverse projection; **E.** UMAP presentation of newly discovered Elastin-VICs, CLDN11-VICs and Proliferative VICs; **F** UMAP visualization of human fetal valves in W22, 23 and 25 (Cui et al., 2019) integrated by CCA; **G.** Feature plots of representative marker genes of each cell type (VICs, VECs, Immune) in Cui et al, 2019. **H.** UMAP visualization of VICs in W22, 23 and 25 (Cui et al., 2019).

### Figure S3

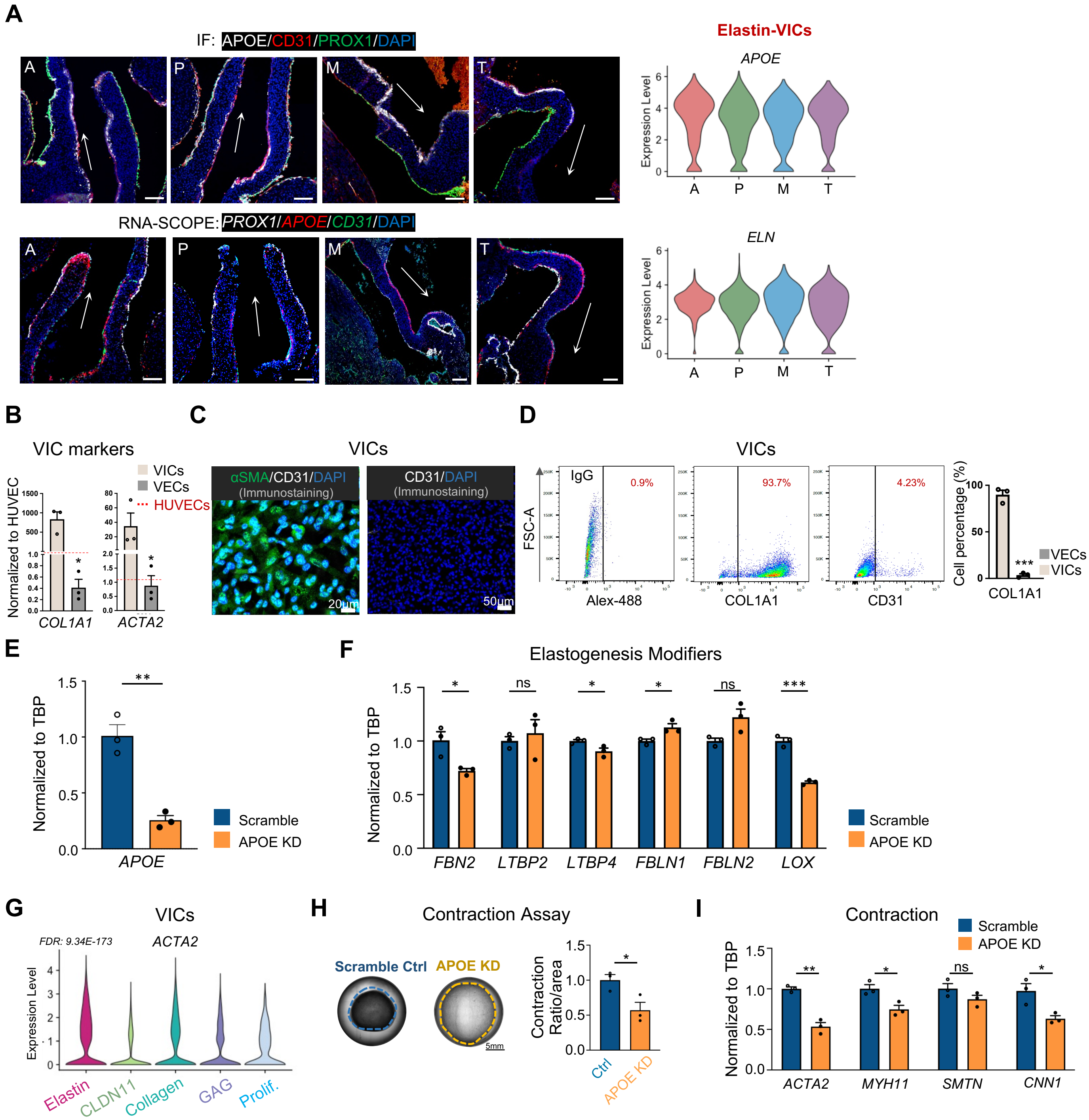


**Figure S3. APOE Expression and Functions in Valve Elastogenesis.**
**A.** Left: Representative immunofluorescence staining and RNA *in situ* hybridization of APOE in four human fetal valves. n=3, Scale bar: 100μm. Arrows represent unidirectional flow directions. Right: Violin plots of *APOE* and *ELN* expression within Elastin-VICs across each valve, A/P/M/T represent Aortic, Pulmonary, Mitral, Tricuspid valve respectively; **B.** qPCR detection of VIC-related genes in cultured VICs, VECs, and HUVECs. Expression levels are normalized to HUVECs; **C.** Immunostaining of αSMA (VICs) and PECAM1 (VECs) on cultured VICs; **D.** Flow cytometry analysis of CD31 (VECs) and COL1A1 (VICs) expression in culture VICs. n=3 biological repeats in each group. **E-F.** qPCR analysis *APOE* (**E**), elastogenesis related genes (**F**) expression in cultured human VICs. n=3 biological repeats; **G.** Violin plots of *ACTA2* expression in VIC subtypes. FDR: Elastin vs. other VIC subclusters; **H.** Contraction assay comparing scramble and APOE KD in VICs. n=3; **I.** qPCR analysis of contraction related genes expression in VICs. n=3 biological repeats. Data shown as the mean ± SEM. *p<0.05, **p<0.01, ***p<0.001, scramble vs APOE KD; Statistics in B, D-F, H-I: Unpaired 2-tailed t-test (2 groups). FDR: False Discovery Rate. HUVECs: Human Umbilical Vein Endothelial Cells.

#### Figure S4

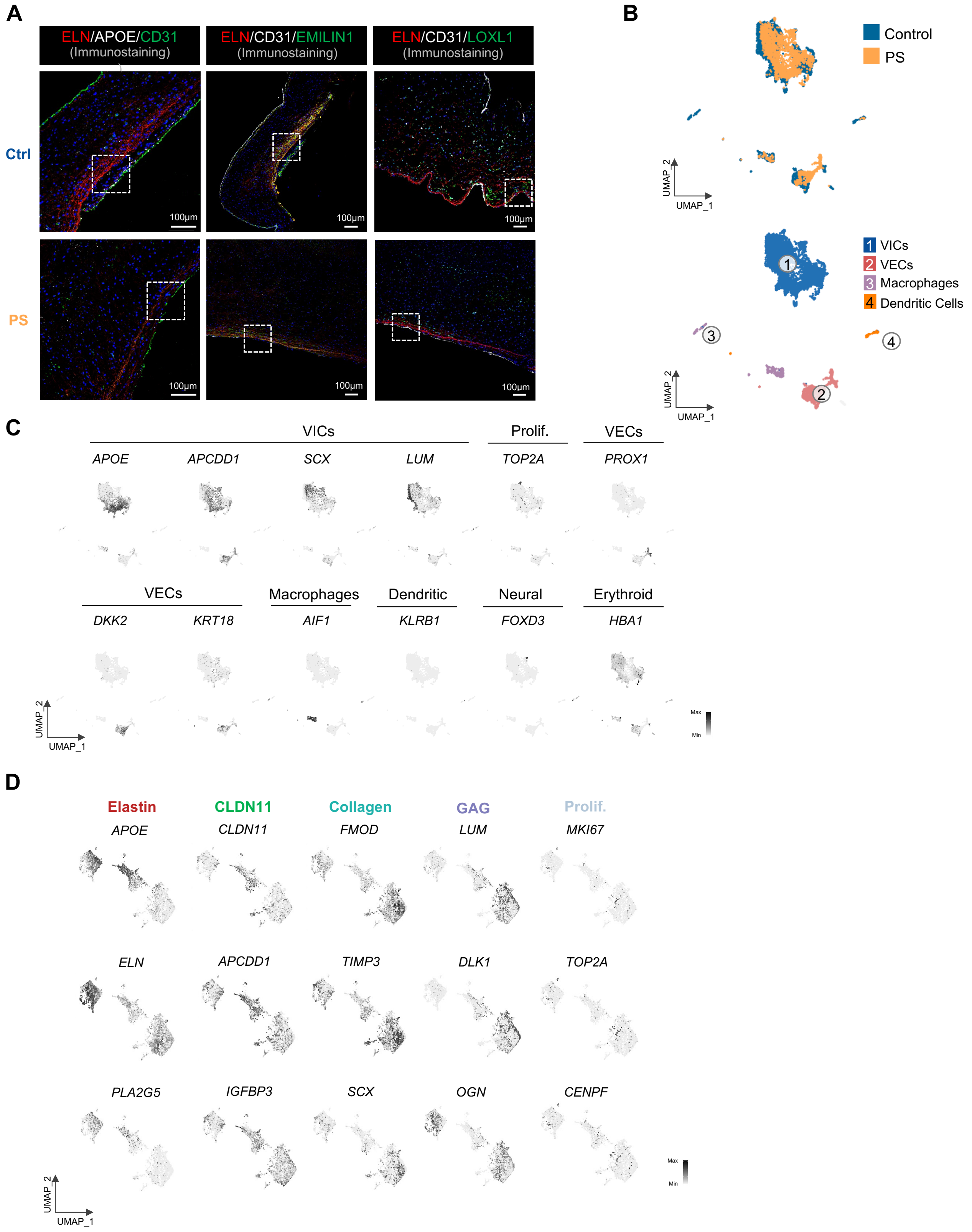


**Figure S4. Overview of Elastin-related Staining and scRNA-seq for PVs with Pulmonary Stenosis.** **A.** Immunofluorescence staining of ELN, APOE, EMILIN1, LOXL1 in pulmonary valves from control and Pulmonary Stenosis (PS). White dashed boxes corresponded to the zoom-in figures shown in Figure 3B. n=3; **B.** UMAP projection of valve cell types (Lower) comparing pulmonary valves (PVs) from healthy control and patient with Pulmonary Stenosis (PS, Upper); **C.** Feature plots of representative marker genes within each cell type and subtypes from B; **D.** Feature plots of representative marker genes of VIC subtypes comparing PVs from healthy control and patient with pulmonary stenosis.

#### Figure S5

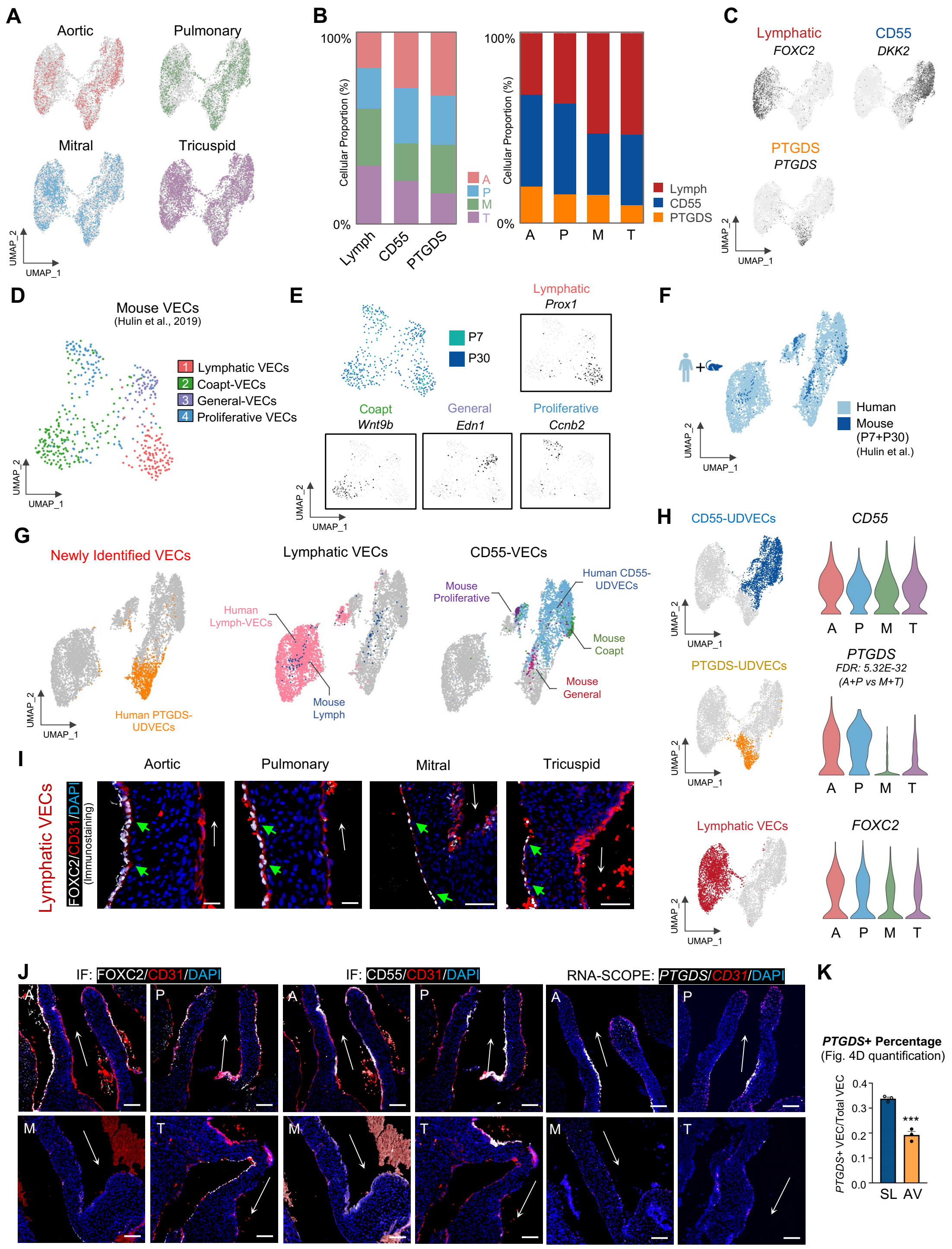


**Figure S5. Overview of VEC Populations from Human and Mouse scRNA-seq and Spatial Location.**
**A.** UMAP demonstrating distribution of four valves within VECs; **B.** Left: Bar graph showing proportions of each valve within each VEC subtype. Right: Bar graph showing proportions of each VEC subtype within each valve; **C.** Feature plots of marker genes from each VEC subtype; **D.** UMAP visualization of mouse VEC subtypes (Hulin et al., 2019); **E.** UMAP visualization of each timepoint, and feature plots of marker genes within each VEC subtype from mouse dataset; **F-G.** UMAP demonstration of the similarity of VEC subtypes between human and mouse VECs through reverse projection; **H.** UMAP visualization of each VEC subtype (Left), violin plot of *CD55*, *PTGDS*, *FOXC2* expression within four valves (Right); **I.** Immunofluorescence staining of FOXC2 in four human fetal valves at W15. n=3, Scale bar: 20μm. White arrows represent unidirectional flow directions; **J.** Immunofluorescence staining of FOXC2, CD55 and RNA *in situ* hybridization of *PTGDS* in four human fetal valves at W15. n=3, Scale bar: 100μm. White arrows represent unidirectional flow directions; **K.** Quantification of *PTGDS*+ VECs from Figure 4D. n=3. Data shown as the mean ± SEM. ***p<0.001. Statistics in K: Unpaired 2-tailed t-test (2 groups). SL: Semilunar valves, AV: Atrioventricular valves.

#### Figure S6

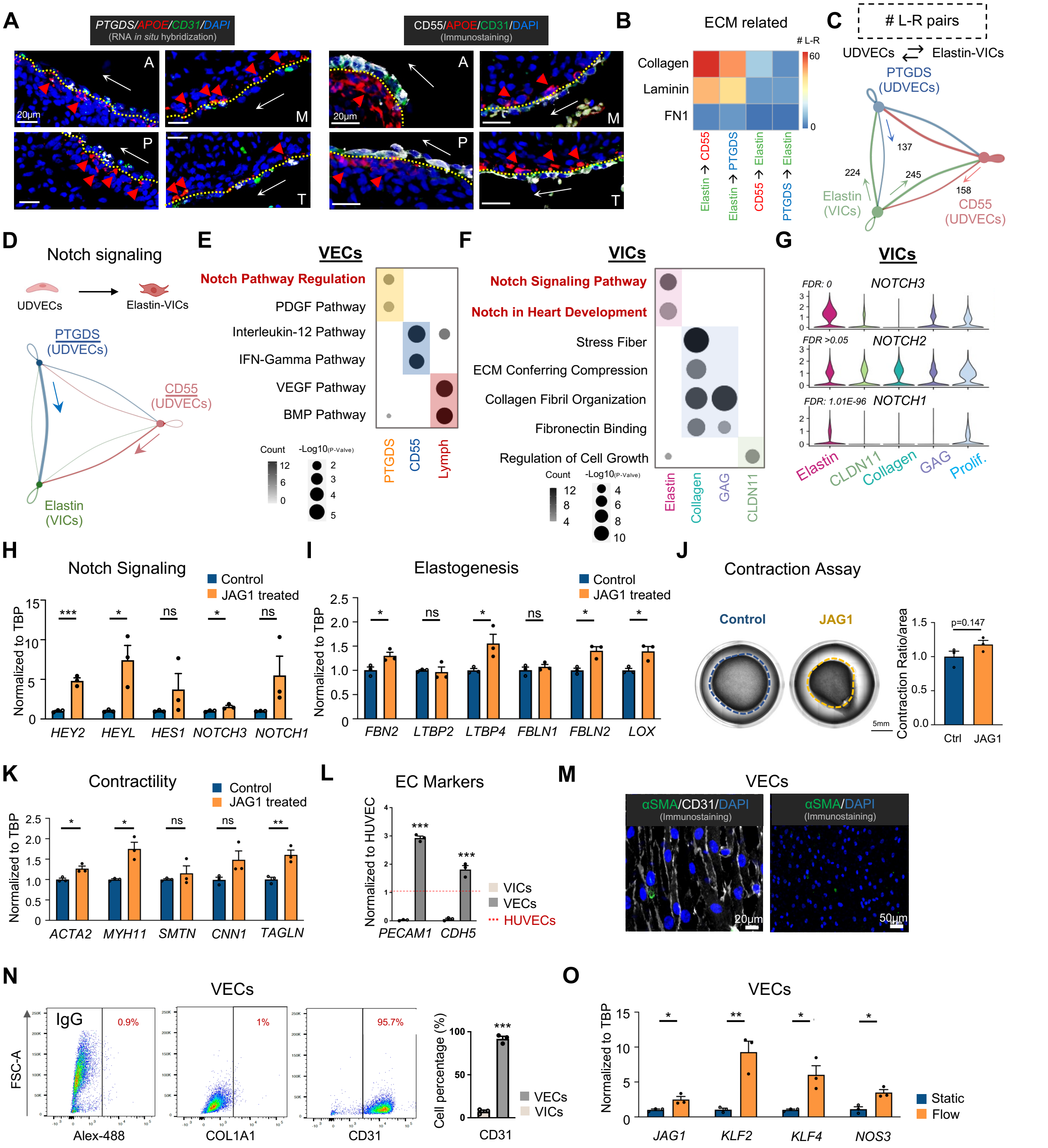


**Figure S6. scRNA-seq Revealed Notch-modulated Elastogenesis between UDVECs and VICs.**
**A.** Co-detection of *APOE* and *PTGDS* through RNA *in situ* hybridization, and APOE with CD55 through immunostaining within four valves. n=3; **B.** Major Extracellular Matrix (ECM) related Ligand-Receptor (L-R) pairs between Elastin-VICs and unidirectional VECs (UDVECs) subtypes; **C.** Total L-R pair numbers between UDVECs and Elastin-VICs; **D.** Notch related signaling patterns from UDVECs to Elastin-VICs; **E-F.** GO enrichment analysis of differential expressed genes within each VEC (**E**) and VIC (**F**) subtype; **G.** Violin plot of Notch receptor expressions in VICs. **H-I.** qPCR analysis of Notch signaling (**H**) and elastogenesis (**I**) related genes in VICs with Jag1 treatment; **J.** Contraction assay of VICs with Jag1 treatment. n=3; **K.** qPCR analysis of contraction related genes in VICs with Jag1 treatment; Jag1: 15ug/ml. **L.** qPCR detection of VEC-related genes in cultured VECs, VICs, and HUVECs. Expression levels are normalized to HUVECs. **M.** Immunostaining of CD31 (VECs) and αSMA (VICs) on cultured VECs; **N.** Flow cytometry analysis of CD31 (VECs) and COL1A1 (VICs) expression in culture VECs. n=3 biological repeats in each group. **O.** qPCR analysis of flow responsive genes and *JAG1* expression in VECs under static or unidirectional flow; Data shown as the mean ± SEM. *p<0.05, **p<0.01, ***p<0.001. Statistics in H-L, N-O: Unpaired 2-tailed t-test (2 groups). ECM: Extracellular Matrix, L-R: Ligand-Receptors

### Figure S7

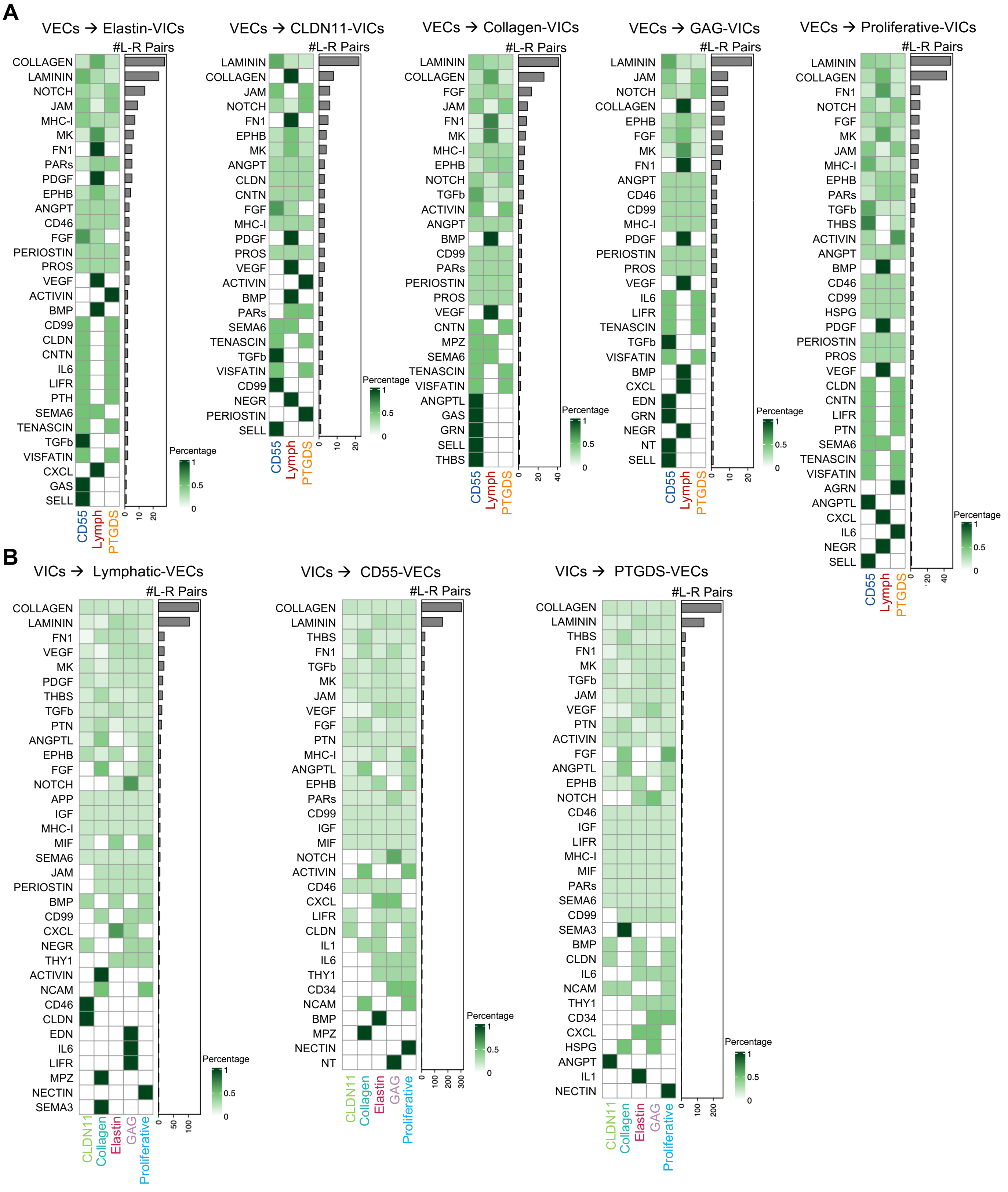


**Figure S7. L-R Pairs Based on scRNA-seq Analysis.**
**A.** Heatmap demonstrating the overview of Ligand-Receptor pair numbers and distribution from VECs to each VIC subtype; **B.** Heatmap demonstrating the overview of Ligand-Receptor pairs numbers and distribution from VICs to each VEC subtype. The number of Ligand-Receptor pairs with probability higher than 0.001 and p value <0.05 were counted for each pathway and visualized by using pheatmap R package.

#### Figure S8

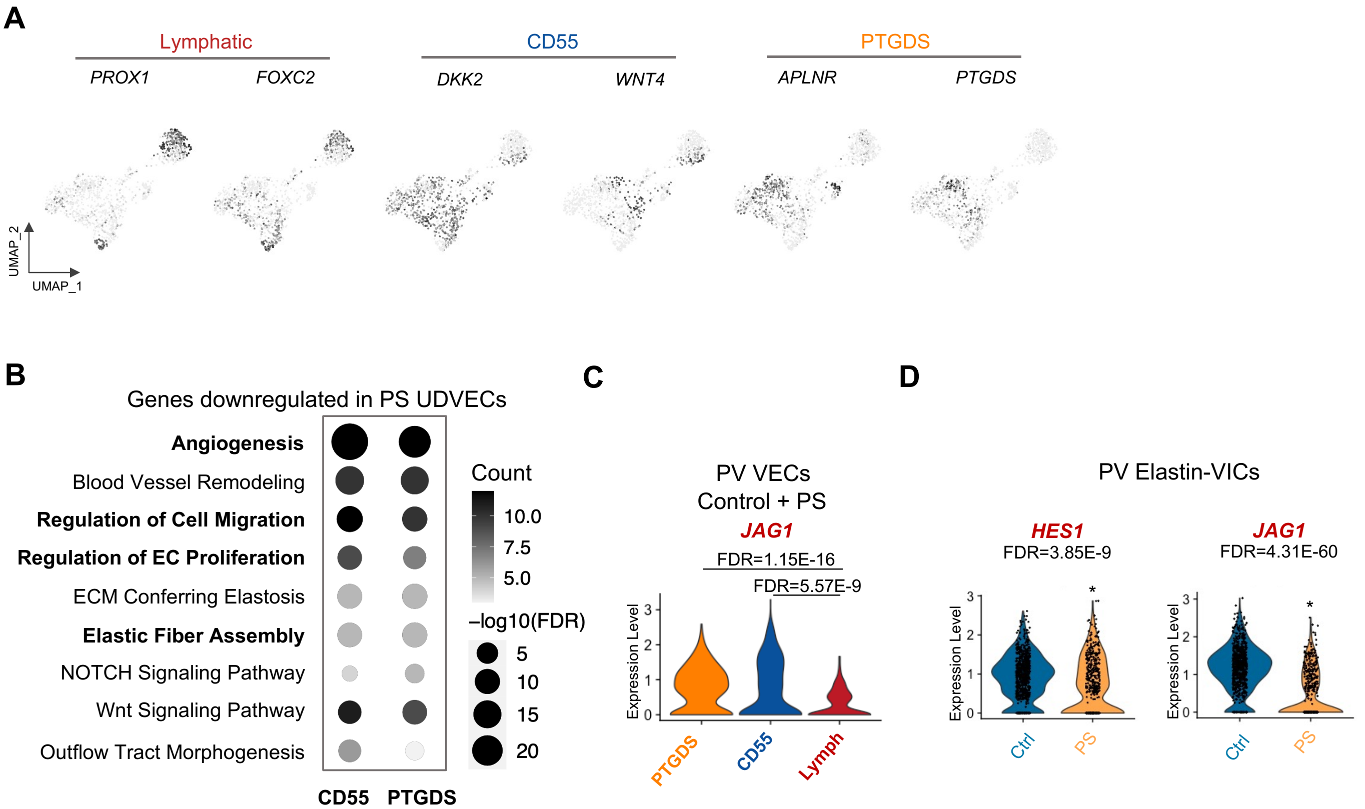


**Figure S8. Overview of VEC scRNA-seq from Pulmonary Valves with Pulmonary Stenosis.**
**A.** Feature plots of markers genes for each VEC subtype combining control and patient with PS; **B.** GO enrichment analysis of genes downregulated in PS unidirectional VECs; **C.** Violin plots comparing *JAG1* expression between three VEC subtypes; **D.** Violin plots comparing *HES1* and *JAG1* expression of control vs. PS in Elastin-VICs.

#### Figure S9

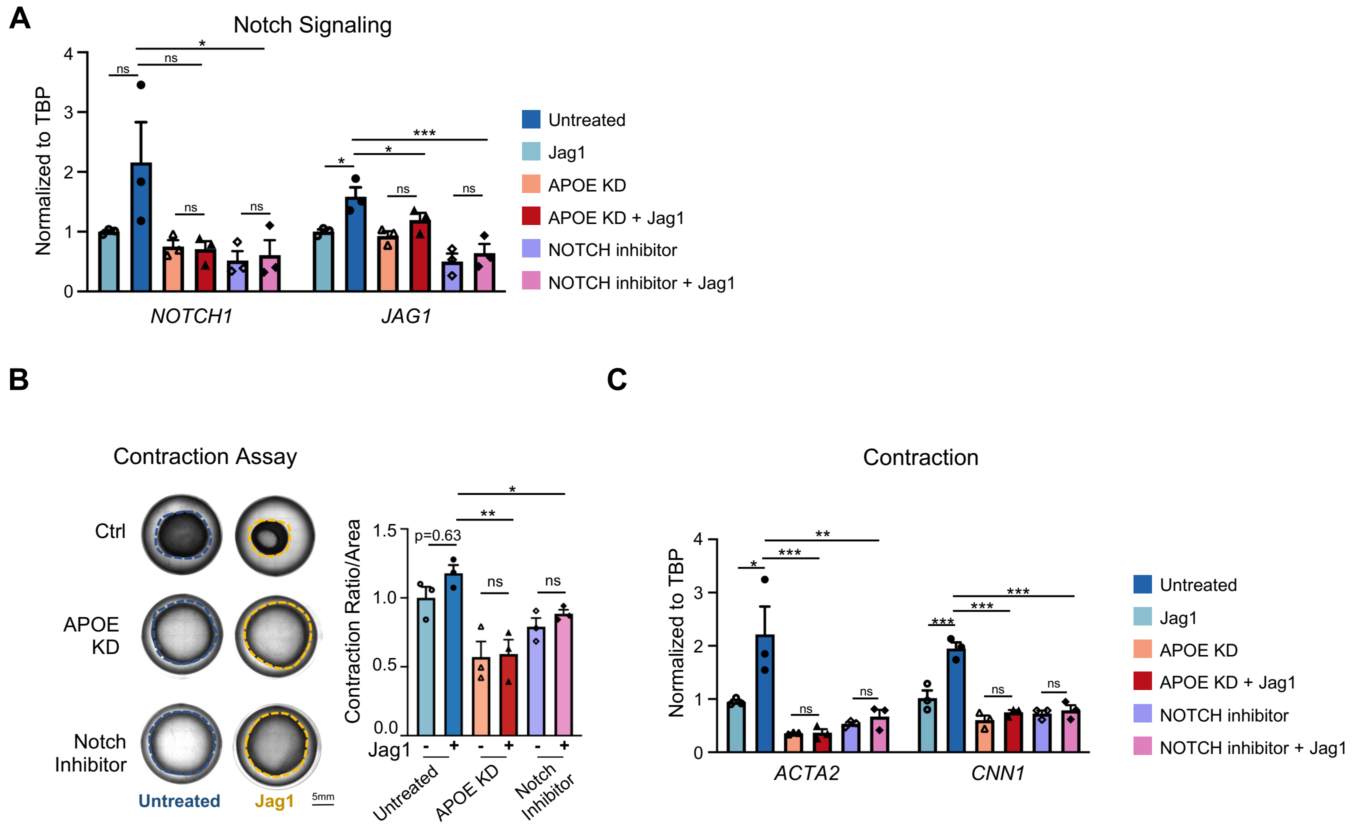


**Figure S9. APOE is Required for Notch Signaling Activation**

**A-C.** VICs were treated with different conditions: Jag1 treatment, APOE knocked down (KD), Notch inhibitor L-685,458 treatment. Jag1: 15ug/ml, L-685,458: 10uM; VICs within each treatment condition were subjected to qPCR analysis and gel contraction assay; **A.** qPCR analysis of *NOTCH1* and *JAG1* in VICs; **B.** Contraction assays of VICs; **C.** qPCR analysis of contraction related genes *ACTA2* and *CNN1* in VICs. Data shown as the mean ± SEM. ns p> 0.05, *p<0.05,**p<0.01,***P<0.001; Statistics in A-C: one-way ANOVA followed by Tukey's test. KD: Knock Down.

#### Reference:


1. Dong Y, Long T, Wang C, Mirando AJ, Chen J, O'Keefe RJ, Hilton MJ. NOTCH-Mediated Maintenance and Expansion of Human Bone Marrow Stromal/Stem Cells: A Technology Designed for Orthopedic Regenerative Medicine. *Stem Cells Transl Med*. 2014;3:1456-1466. doi: 10.5966/sctm.2014-0034
2. Miyagawa K, Shi M, Chen PI, Hennigs JK, Zhao Z, Wang M, Li CG, Saito T, Taylor S, Sa S, et al. Smooth Muscle Contact Drives Endothelial Regeneration by BMPR2-Notch1-Mediated Metabolic and Epigenetic Changes. *Circ Res*. 2019;124:211-224. doi: 10.1161/CIRCRESAHA.118.313374
3. Warboys CM, Ghim M, Weinberg PD. Understanding mechanobiology in cultured endothelium: A review of the orbital shaker method. *Atherosclerosis*. 2019;285:170-177. doi: 10.1016/j.atherosclerosis.2019.04.210
4. Gu M, Shao NY, Sa S, Li D, Termglinchan V, Ameen M, Karakikes I, Sosa G, Grubert F, Lee J, et al. Patient-Specific iPSC-Derived Endothelial Cells Uncover Pathways that Protect against Pulmonary Hypertension in BMPR2 Mutation Carriers. *Cell Stem Cell*. 2017;20:490-504 e495. doi: 10.1016/j.stem.2016.08.019
5. Tran HTN, Ang KS, Chevrier M, Zhang X, Lee NYS, Goh M, Chen J. A benchmark of batch-effect correction methods for single-cell RNA sequencing data. *Genome Biol*. 2020;21:12. doi: 10.1186/s13059-019-1850-9
6. Hafemeister C, Satija R. Normalization and variance stabilization of single-cell RNA-seq data using regularized negative binomial regression. *Genome Biol*. 2019;20:296. doi: 10.1186/s13059-019-1874-1
7. Xie C, Mao X, Huang J, Ding Y, Wu J, Dong S, Kong L, Gao G, Li CY, Wei L. KOBAS 2.0: a web server for annotation and identification of enriched pathways and diseases. *Nucleic Acids Res*. 2011;39:W316-322. doi: 10.1093/nar/gkr483
8. Jin S, Guerrero-Juarez CF, Zhang L, Chang I, Ramos R, Kuan CH, Myung P, Plikus MV, Nie Q. Inference and analysis of cell-cell communication using CellChat. *Nat Commun*. 2021;12:1088. doi: 10.1038/s41467-021-21246-9
